# The CGG triplet repeat binding protein 1 counteracts DNA secondary structure-induced transcription-replication conflicts

**DOI:** 10.1101/2023.03.09.531843

**Authors:** Henning Ummethum, Maxime Lalonde, Marcel Werner, Manuel Trauner, Anna Chanou, Matthias Weiβ, Clare S. K. Lee, Elisabeth Kruse, Andreas Ettinger, Stephan Hamperl

## Abstract

Short tracts of trinucleotide repeats with less than 10 repeats are found frequently throughout the genome without any apparent negative impact on DNA replication fork progression or transcription elongation. CGG binding protein 1 (CGGBP1) binds to CGG triplet repeats and has been implicated in multiple cellular processes such as transcription, replication and DNA damage. Here, we show that CGGBP1 binds to human gene promoter sites prone to G-quadruplex and R-loop secondary structure formation. Altering CGGBP1 levels results in the accumulation of R-loops and causes a defect in transcriptional elongation by RNA polymerase II, which subsequently leads to replication fork stalling and transcription-replication conflicts. Together, our work shows that short trinucleotide repeats are a source of genome-destabilizing secondary structures and cells rely on specific DNA-binding factors to maintain proper transcription and replication progression at short trinucleotide repeats.

## Introduction

Secondary DNA structures arising in the genome as a result of transcription and replication can be a threat to genome stability (Aguilera and Gaillard, 2014; Chang et al., 2017; French, 1992; Fu et al., 2011; Hamperl et al., 2017; Promonet et al., 2020; Sankar et al., 2016; Srivatsan et al., 2010; Tuduri et al., 2009; Urban et al., 2016). Both transcription and replication proceed by unwinding the template DNA, thereby creating single-stranded DNA (ssDNA) and potentially allowing stable secondary DNA structures such as hairpins, cruciforms, triplexes (H DNA) and G L quadruplexes (G4s) to form. Tracts of repetitive sequences such as CGG-trinucleotide repeats L are particularly prone to secondary structure formation. The formation of secondary structures can lead to the expansion of CGG repeats, for example in the 5’ untranslated region (UTR) of the human Fragile X Messenger Ribonucleoprotein 1 (FMR1) gene (Zhao and Usdin, 2016), which is observed in the human neurodegenerative disease Fragile X syndrome (FXS) (Verkerk et al., 1991). When the CGG expansion exceeds 200 repeats they become hypermethylated, causing transcriptional silencing of the FMR1 gene. This results in the loss of its encoded protein FMRP, which is a selective RNA-binding protein implicated in many cellular functions such as regulating dendritic mRNA transport, local protein synthesis at synapses (Feng et al., 1997; Fu et al., 1991; Pieretti et al., 1991) among many others (reviewed in Davis and Broadie, 2017).

Disease-length repetitive CGG-tracts are prone to form R loops (Groh et al., 2014; Loomis et al., 2014), These RNA:DNA hybrids form when nascent transcripts reanneal to their template DNA, displacing the non-template strand as ssDNA. R-loop formation occurs co-transcriptionally and has been proposed to impact both transcription and replication by reducing RNAPII elongation and hindering replication fork progression. Despite the clinically relevant example of CGG-trinucleotide repeat expansion at the FMR1 gene, short trinucleotide repeat tracts with n < 10 are extensively scattered across the entire genome (Clark et al., 2006; Willems et al., 2014). However, these short repeats are not known to impact transcription and/or replication, despite the ability of even (CGG)5 oligonucleotides to form a stable G-quadruplex structure at physiological pH *in vitro* (Fry and Loeb, 1994). Further, these “normal” repeats do not significantly induce R loops compared to longer repeats associated with FXS and are not associated with delay of RNAPII or transcriptional silencing (Groh et al., 2014). However, whether such short repeats are capable to form secondary structures *in vivo* or whether and how the formation of harmful G4 and R-loops at those repeats is regulated, remains unclear.

The CGG binding protein 1 (CGGBP1) was identified as a short CGG triplet repeat binding protein *in vitro* (Deissler et al., 1996, 1997; Richards et al., 1993). CGGBP1 is a 20kDa protein with a nuclear localization signal and a predicted C2H2-type Zn finger DNA-binding domain (Müller-Hartmann et al., 2000; Singh and Westermark, 2015). Electrophoretic mobility shift assays with CGG-repeat containing oligonucleotides confirmed strong affinity to CGG-repeat sequences *in vitro* (Deissler et al., 1996; Müller-Hartmann et al., 2000), which has also been verified *in vivo* at selected loci, such as the CGG repeats in the 5’ UTR of the FMR1 gene (Goracci et al., 2016). Overexpression of CGGBP1 promotes the binding onto the 5’ UTR of the FMR1 gene leading to transcriptional repression (Müller-Hartmann et al., 2000), suggesting that an excess of CGGBP1-bound repeats may have a dominant negative effect on transcription elongation. On the other hand, CGGBP1 knockdown did not affect either FMR1 transcription activity in transcriptionally active alleles or CGG expansion stability in cell lines with expanded FMR1 CGG alleles (Goracci et al., 2016). Such a gene- and context-specific effect of CGGBP1 levels on transcription regulation was also observed at other gene promoters. For example, CGGBP1 binding at the HSF1 promoter is required both for driving basal levels of transcription as well as for repressing excessive levels of expression that are permitted only after heat shock induction (Singh et al., 2009). One possibility is that CGGBP1 has a direct regulatory effect on RNAPII elongation during passage of the transcription complex through CGGBP1-bound repeats, but the molecular details of such differential interactions have not been investigated.

Here we first profiled CGGBP1 binding genome-wide and identified its target binding sites. We find that CGGBP1 is strongly enriched at short CGG tandem repeats, in particular at transcribed gene promoter regions. Downregulation of CGGBP1 in human U-2OS cells results in a global increase of transcriptional activity. Likewise, overexpression of CGGBP1 decreases global RNA synthesis rates, suggesting that CGGBP1 functions as a global transcriptional repressor, as previously reported (Singh and Westermark, 2015). Interestingly, a subset of RNAPII genes containing direct binding sites of CGGBP1 require basal levels of CGGBP1 for efficient transcription elongation. Consistently, loss of CGGBP1 results in more chromatin-associated RNAPII in S-phase, thereby causing replication fork impairment and an elevated frequency of transcription-replication conflicts (TRCs). Using an episomal system, we provide direct evidence that CGGBP1 can prevent the formation of transcription-impairing and genome-destabilizing R-loops. Together, these observations suggest that CGGBP1 has key functions in the regulation of transcription without provoking excessive formation of DNA secondary structures such as G4s and R-loops over short CGG tandem repeats that would otherwise become genome-destabilizing transcription and replication impediments.

## Results

### Global profiling of CGGBP1 binding sites in the human genome

To characterize the genome-wide binding targets of CGGBP1, we first took advantage of a publicly available ENCODE CGGBP1 Chromatin Immunoprecipitation-Sequencing (ChIP-Seq) dataset in human K562 cells (The ENCODE Project Consortium, 2012). We identified a total of 3459 CGGBP1 binding sites across two biological replicates with high confidence over the untagged control (**Figures 1A-C**). Strikingly, the majority of identified ChIP-Seq peaks (∼ 90%) mapped onto non-repetitive euchromatic regions with high gene density, with a strong over-representation at RNAPII-dependent promoter regions. In fact, ∼73% of the peaks were found within 1kb genomic distance to gene promoters and more than 80% of the detected CGGBP1 peaks resided within 3kb distance to the nearest transcription start site (TSS) (**Figures 1A-B**). A small fraction of target genes, such as MALAT1 and RPL23A/TLCD1, displayed CGGBP1 peaks over the whole gene body with a preference for intronic sequences (**Figures 1A-B**). About 10% of all peaks occurred at intergenic regions distal from annotated TSSs and gene body regions (**Figure 1B**). This suggests that the majority of CGGBP1 binding events occur within genic regions with a strong preference towards gene promoters of RNAPII-transcribed genes.

**Figure 1.**
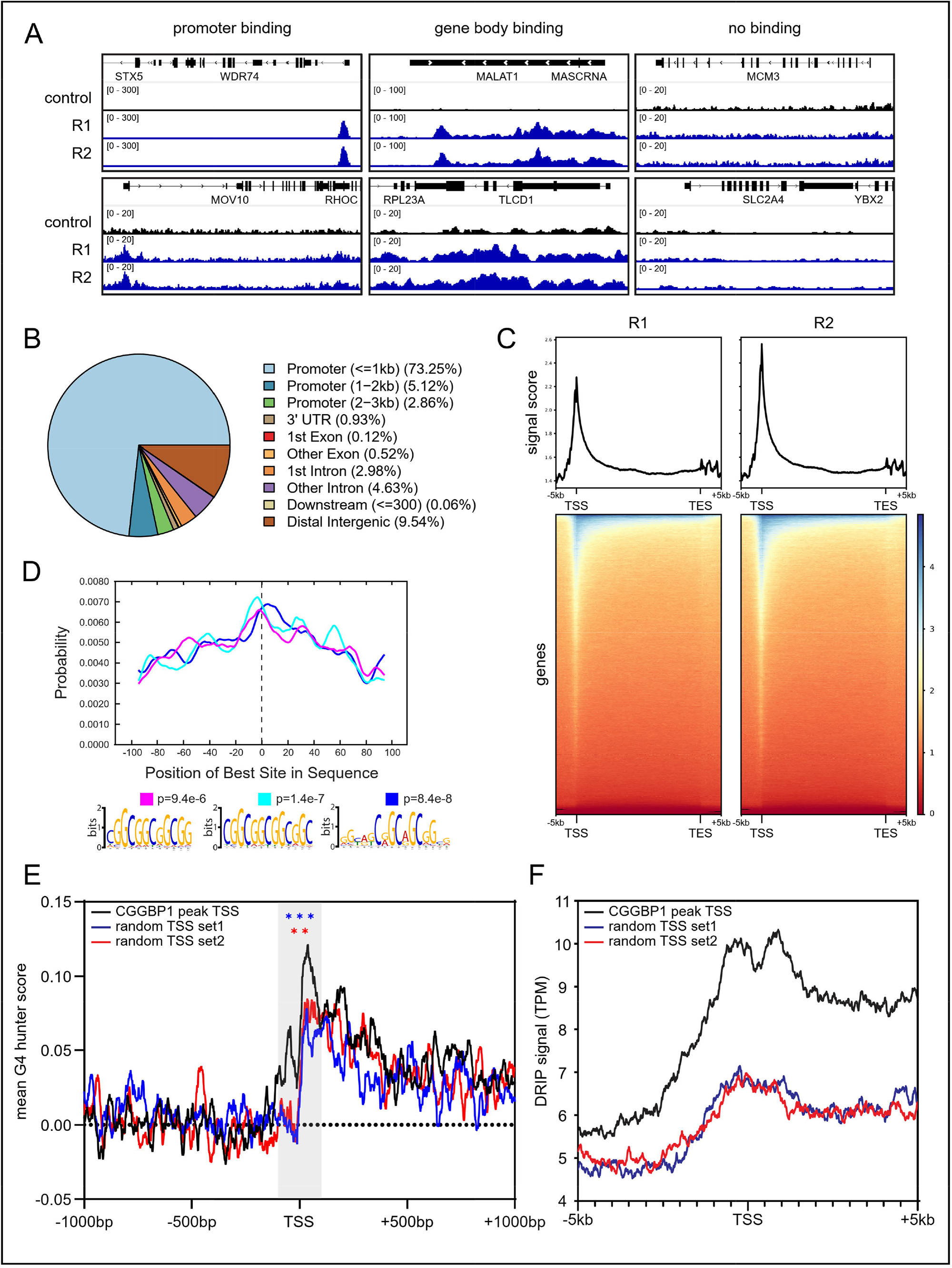
Global profiling of CGGBP1 binding sites in the human genome. A) Example tracks of CGGBP1 ChIP-seq signal in K562 cells at different genes of two replicates compared to the control. B) Pie chart showing the distribution of CGGBP1 ChIP-seq peak annotation relative to the position in the genome. C) Metagene plots and underlying heatmaps of CGGBP1 ChIP-seq signal of two replicates relative to the control across all annotated transcripts. D) Motif probability graph and DNA sequence logos of the most significant hits found in the CGGBP1 ChIP-seq peak sequences. E) Summary plot of predicted G4 quadruplex formation scores at either TSSs with a CGGBP1 binding peak in proximity or two random sets of TSSs. Statistical significance was calculated using two tailed student’s t-test on the medians of 10 bp windows at the TSS +/-100 bp indicated by the grey box. Significance was tested against both controls in individual t-tests indicated by the color of the stars. F) Summary plot of DRIP-seq signal at either TSSs with a CGGBP1 binding peak in proximity or two random sets of TSSs.

As expected, CGGBP1-positive regions were enriched for short CGG repeats. Motif identification using MEME revealed three motifs containing up to four consecutive CGG repeats (**Figure 1D**). Consistent with the *in vitro* binding preference of CGGBP1 towards CGG repeats (Deissler et al., 1996, 1997; Richards et al., 1993), these findings confirmed that CGGBP1 preferentially binds short tracts of CGG-repeats *in vivo*, a motif that is enriched at RNAPII-dependent promoters (Sawaya et al., 2013). Due to this inherent DNA sequence bias, the TSSs containing a CGGBP1 peak showed an elevated level of GC-content and GC-skew in comparison to two matched sets of randomly shuffled control TSSs without CGGBP1 peaks (**Figures S1A-B**). Because both GC-content and GC-skew are positively correlated with the formation of G4 and R-loops (Gellert et al., 1962; Lam et al., 2013; Sen and Gilbert, 1988; Shrestha et al., 2014), we next asked whether CGGBP1-bound sites show increased levels of G4 or R-loops. We used G4 Hunter to predict G4 structure formation in a region 1kb upstream and downstream of the TSS (Bedrat et al., 2016). We found that CGGBP1 bound TSSs have a higher probability to form G4s within -/+100bp of the TSS compared to two randomly permutated controls with an equal number of TSSs (**Figure 1E**). Analysis of a DRIP-Seq dataset from U2OS cells also indicated a strong increase in R-loop signal at CGGBP1-bound TSSs compared to control TSSs (**Figure 1F**). Together, these data suggest that CGGBP1 binding sites in the genome coincide with sites of short CGG-trinucleotide repeats at RNAPII promoters and show a high propensity to form G4 and R-loops.

### Altering cellular CGGBP1 levels leads to global transcriptional changes

Given its localisation to gene promoters, we next asked whether CGGBP1 can regulate RNAPII transcription. We first performed siRNA-mediated knockdown of CGGBP1 using a pool of four independent CGGBP1-targeting siRNAs, which led to efficient depletion of CGGBP1 after 24h of transfection (**Figure 2A**). Prolonged knockdown of CGGBP1 for 48h and 72h resulted in an increased accumulation of cells in G1-phase (**Figure 2B** **and Figure S2A**), consistent with previous studies (Singh et al., 2011). Thus, a basal expression level of CGGBP1 is important for cell cycle progression.

**Figure 2.**
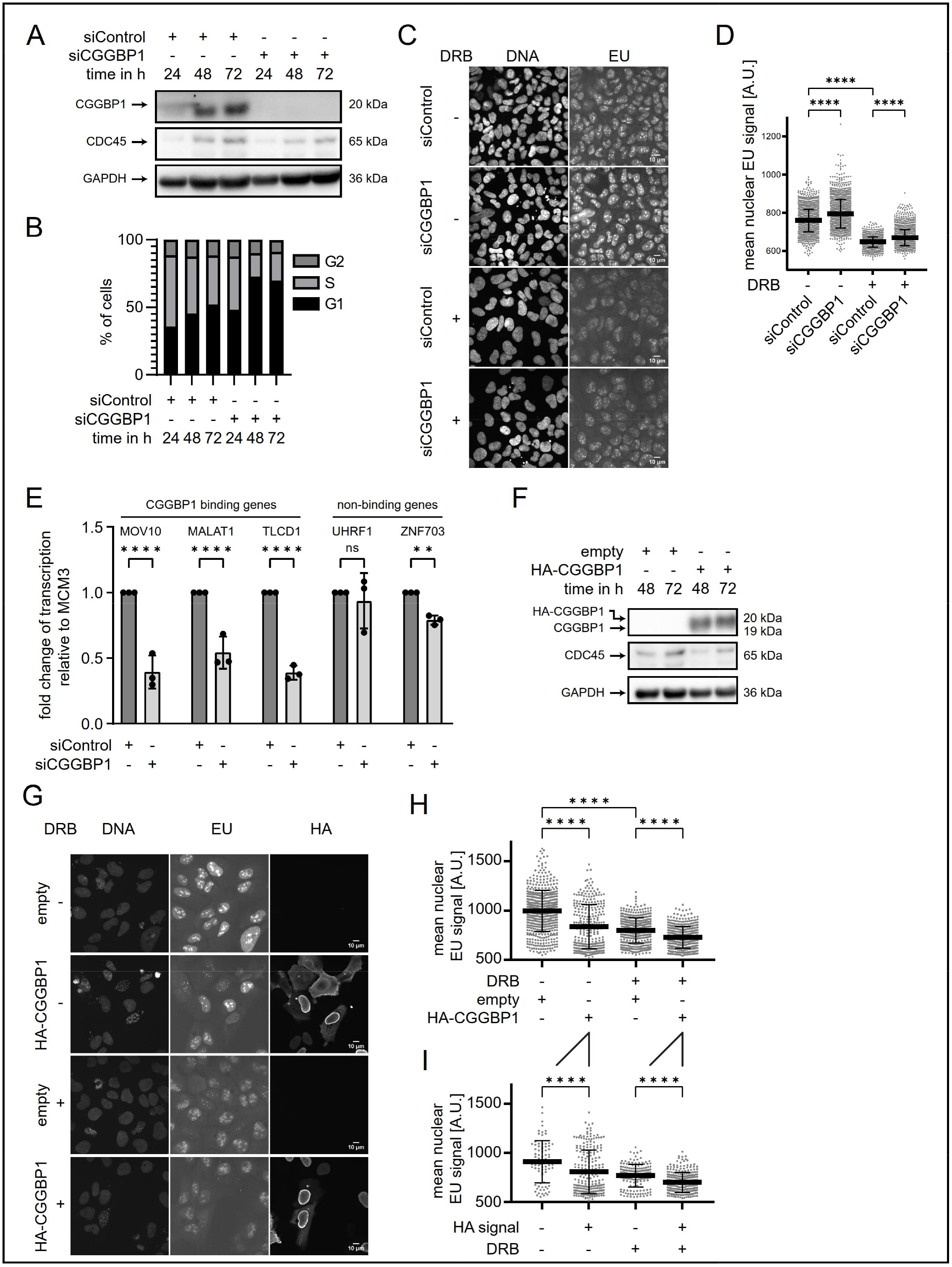
Altering cellular CGGBP1 levels leads to global transcriptional changes. A) Western Blot of U2OS total cell extracts after 24, 48 and 72 h of CGGBP1 knockdown compared to control siRNA. CDC45 and GAPDH are loading controls. N=1. B) Quantification of cell cycle distribution in BrdU cell cycle analysis by flow cytometry upon treatment of U2OS cells with siControl or siCGGBP1 for 24, 48 and 72 h (flow cytometry profiles in **Figure S2A**). N=1. C) Example immunofluorescence (IF) images of EU incorporation in U2OS cells upon 72 h CGGBP1 knockdown compared to control siRNA. For transcriptional inhibition, 100 µM DRB was added 2 h before fixation. N=1. D) Quantification of mean nuclear EU signal in all fields similar to C). For transcriptional inhibition, 100 µM DRB was added 2 h before fixation. Data are represented as mean ± standard deviation. Statistical significance was calculated using ordinary one-way ANOVA. N=1. E) Gene expression of selected examples measured by RT-qPCR of cDNA from U2OS Tet-ON pHU43 clone 2 (see Figure 6) treated with siControl or siCGGBP1 for 72 h and no DOX. Shown is the fold change relative to MCM3 and normalized to siControl. Statistical significance was calculated using ordinary one-way ANOVA. N=3. F) Western Blot of HA-CGGBP1 overexpression in U2OS cells. Total cell extracts were taken 48 or 72 h after transfection of the overexpression plasmid. CDC45 and GAPDH are loading controls. WT CGGBP1 is not visible here due to the high amount of HA-CGGBP1. N=1. G) Example IF images of EU incorporation and HA signal in U2OS cells upon HA-CGGBP1 overexpression compared to an empty overexpression plasmid control. For transcriptional inhibition, 100 µM DRB was added 2 h before fixation. N=1. H) Quantification of mean nuclear EU signal in all fields similar to G). Data are represented as mean ± standard deviation. Statistical significance was calculated using ordinary one-way ANOVA. N=1. I) Quantification of mean nuclear EU signal in all fields similar to G). Only the overexpression condition is shown with cells split into HA+ or HA-. Statistical significance was calculated using ordinary one-way ANOVA. N=1.

To investigate whether CGGBP1 regulates transcription, we pulse-labelled cells with the uridine analogue 5-ethynyluridine (EU), which is incorporated into newly synthesized RNA (Jao and Salic, 2008). Interestingly, we observed a significant increase in transcriptional activity in CGGBP1-depleted cells compared to control siRNA cells (**Figure 2C-D**). Importantly, treatment with transcription elongation inhibitor 5,6-Dichlorobenzimidazole 1-β-D-ribofuranoside (DRB), a specific inhibitor of RNAPII Serin-2 phosphorylation (Bensaude, 2011; Zandomeni et al., 1982), significantly reduced nuclear EU signal in both siControl and siCGGBP1 cells. However, CGGBP1 depleted cells maintained an elevated level of EU incorporation under conditions of RNAPII transcription inhibition. These data suggest that the increase in EU signal may reflect RNAPI or RNAPIII activity (**Figure 2C-D**).

To better address the relationship between CGGBP1 binding at RNAPII promoters and transcriptional activity, we measured the expression level of three genes that contain CGGBP1 binding sites (MOV10, MALAT1 and TLCD1, **Figure 1A**) and two genes without CGGBP1 binding (UHRF1 and ZNF703) by RT-qPCR. For all three CGGBP1-bound target genes, we observed a strong reduction in mRNA levels in CGGBP1-depleted cells to ∼40-50% compared to the control siRNA cells, whereas non-CGGBP1-bound genes were only mildly (ZNF703) or not affected (UHRF1) (**Figure 2E**). This suggests that CGGBP1 positively regulates its target genes. Furthermore, this implies that the global increase in transcriptional activity upon depletion of CGGBP1 (**Figure 2C-D**) is not related to direct CGGBP1-bound RNAPII target genes.

Next, we wanted to understand how overexpression of CGGBP1 globally affects nascent transcription levels. To this end, we transiently overexpressed HA-tagged CGGBP1, which led to high levels of CGGBP1 as determined using an HA-antibody (**Figure 2F**). We then determined EU incorporation rates in HA (-) and HA (+) cells with and without RNAPII transcription inhibition by DRB treatment. CGGBP1-overexpressing cells showed a significant decrease in nascent transcription levels, suggesting that increased CGGBP1 levels negatively affect global RNA synthesis (**Figures 2G-H**). This decrease of EU incorporation of HA (+) versus HA (-) cells was also observed in the presence of DRB, again suggesting that this effect was not limited to RNAPII transcription. Importantly, the decrease in transcriptional activity was only observed in transfected HA (+) cells (**Figure 2I**), suggesting that the effects on transcriptional activity were caused by CGGBP1. Together, these data suggest that high levels of CGGBP1 lead to global transcriptional repression. Thus, a balanced level of cellular CGGBP1 levels appears to be critical for correct nuclear transcription rates (including all three nuclear RNA Polymerases), as knockdown and overexpression of CGGBP1 positively and negatively impacts transcription on a global level, respectively.

### Altering cellular CGGBP1 levels impacts the level of chromatin-bound RNAPII complexes

As gene promoters are the major regulatory sites for RNAPII loading and transcription initiation and represent the primary target sites of CGGBP1 binding (**Figure 1**), we asked if and how CGGBP1 binding at promoters affects RNAPII occupancy on chromatin. First, we performed chromatin fractionation and measured the levels of total chromatin-bound RNAPII versus elongating form of RNAPII phsopho-Ser2 (RNAPII-pS2) by Western blot (**Figure S3A-B**). We observed a ∼1.5-fold increase in the amount of RNAPII-pS2 on chromatin in CGGBP1-depleted cells compared to control knockdown cells, suggesting that CGGBP1 has a role in regulating this ratio of gene-body associated compared to total chromatin-bound RNAPII complexes (**Figure S3A-B**). As expected, DRB treatment decreased the ratio of RNAPII-pS2 levels over total RNAPII. Triptolide, an XPB/TFIIH inhibitor (Leuenroth and Crews, 2008; Titov et al., 2011), induced rapid proteasomal degradation of both RNAPII-pS2 and total RNAPII complexes, confirming the specificity of the used antibodies (**Figures S3A-B**).

To confirm this result with a more quantitative method and discriminate between different cell cycle stages, we performed immunofluorescence of pre-extracted cells to determine the levels of chromatin-bound total RNAPII and RNAPII-pS2 per cell. We also included pulse labelling of cells with 5-Ethynyl-2’-deoxyuridine (EdU) to discriminate between S-phase and non-S-phase cells (**Figure 3A-C** **and Figures S3C-F**). The S-phase population was reduced upon CGGBP1 depletion, confirming the flow cytometry results (**Figure 3C** **and** **Figure 2B**). CGGBP1 knockdown led to a small but significant increase in both chromatin-bound RNAPII and RNAPII-pS2 signals (**Figures S3C-F**). Transcription inhibition by DRB resulted in a significant decrease in nuclear intensities, although CGGBP1-depleted cells always retained higher levels compared to control siRNA cells (**Figure S3C-F**). Interestingly, calculating the per-cell ratio of RNAPII-pS2 versus total RNAPII in EdU (-) and EdU (+) cells showed that the increase in RNAPII-pS2 signal in CGGBP1-depleted cells was mainly derived from S-phase cells (**Figure 3B**), suggesting that CGGBP1 mitigates the accumulation of chromatin-bound RNAPII-pS2 during active DNA replication. Importantly, transcription inhibition by DRB rescued this effect in CGGBP1-depleted S-phase cells (**Figure 3B**), supporting the notion that this CGGBP1-induced stalling of RNAPII is transcription-dependent.

**Figure 3.**
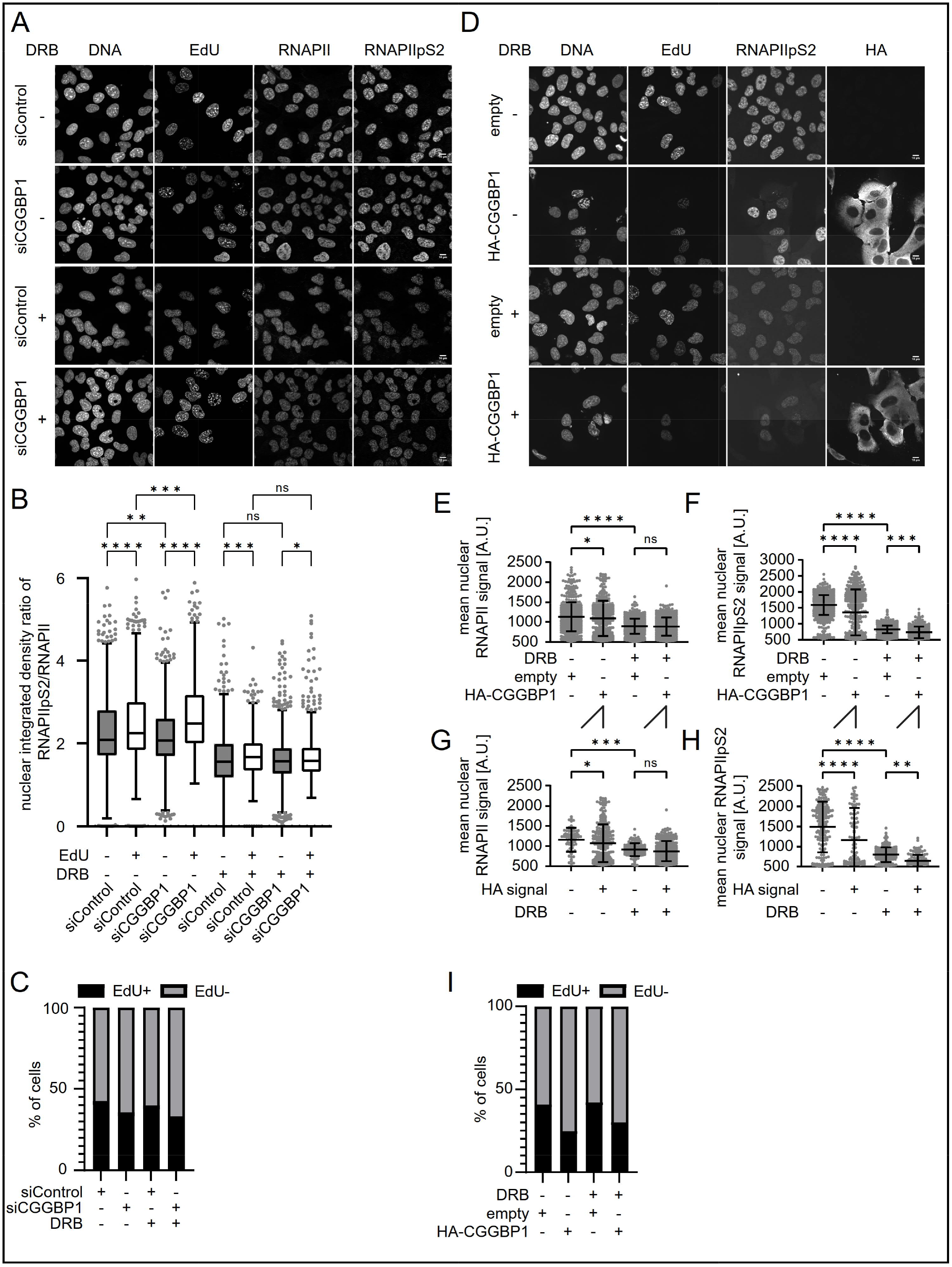
Altering cellular CGGBP1 levels impacts the level of chromatin-bound RNAPII complexes. A) Example IF images of total RNAPII, RNAPII pS2 and EdU incorporation upon treatment of U2OS cells with siControl or siCGGBP1 for 72 h. For transcriptional inhibition, 100 µM DRB or 1 µM triptolide were added 2 h before fixation. N=1. B) Box plot of ratios between RNAPII pS2 and total RNAPII mean integrated nuclear signal in the same cell calculated for EdU+ and EdU-cells in each condition of A). Statistical significance was calculated using ordinary one-way ANOVA. N=1. Further quantifications in **Figure S3**. C) Bar graph showing the percentage of EdU+ and EdU-cells in all fields similar to A). N=1. D) Example IF images of RNAPII pS2 and EdU incorporation in U2OS cells upon HA-CGGBP1 overexpression compared to an empty overexpression plasmid control. For transcriptional inhibition, 100 µM DRB was added 2 h before fixation. N=1. E) Quantification of mean nuclear total RNAPII signal in in all conditions of D). Data are represented as mean ± standard deviation. Statistical significance was calculated using ordinary one-way ANOVA. N=1. Further quantifications in Figure S3. F) Quantification of mean nuclear RNAPII-pS2 signal in all fields similar to D). Data are represented as mean ± standard deviation. Statistical significance was calculated using ordinary one-way ANOVA. N=1. G) Quantification of mean nuclear total RNAPII signal in all conditions of D). Only the overexpression condition is shown with cells split into HA+ or HA-. Data are represented as mean ± standard deviation. Statistical significance was calculated using ordinary one-way ANOVA. N=1. H) Quantification of mean nuclear RNAPII-pS2 signal in all fields similar to D). Only the overexpression condition is shown with cells split into HA+ or HA-. Data are represented as mean ± standard deviation. Statistical significance was calculated using ordinary one-way ANOVA. N=1. I) Bar graph showing the percentage of EdU+ and EdU-cells in all fields similar to D). N=1.

To analyse more specifically the effect of CGGBP1 overexpression on RNAPII chromatin binding, we performed a similar immunofluorescence assay against total RNAPII and RNAPII-pS2 in cells overexpressing HA-CGGBP1 and control cells (**Figure 3D-I** **and Figure S3G-H**). In addition, we included pulse-labelling with EdU and observed a significant decrease of EdU-positive cells in CGGBP1-overexpressing cells (**Figure 3I**), similar to what we observed in CGGBP1 knockdown cells (**Figure 3C**). Interestingly, chromatin-bound RNAPII-pS2 levels dramatically decreased in CGGBP1-overexpressing cells, whereas only a mild effect was observed for total RNAPII (**Figures 3E-F**), suggesting that high levels of CGGBP1 impair RNAPII-pS2. Consistently, splitting the HA-CGGBP1 transfected cell population into HA (-) and HA (+) cells revealed that this decrease was only observed in transfected HA (+) cells (**Figure 3G-H**), providing strong support that this effect was directly correlated with CGGBP1 levels and not due to secondary effects from transfecting different constructs. Since RNAPII-pS2 is a mark of elongating RNAPII, these data are consistent with the preferential enrichment of CGGBP1 binding sites at promoter regions and implies that CGG repeats decorated with high levels of CGGBP1 impair transcription initiation and/or promoter escape/transcription elongation.

### CGGBP1 levels are important to mitigate transcription-replication interference

Given that the accumulation of RNAPII-pS2 in CGGBP1-depleted cells occurs preferentially during the S-phase (**Figure 3B**), we wondered whether CGGBP1’s transcription regulatory role was associated with a putative role in managing transcription-replication conflicts (TRCs). To evaluate TRC levels, proximity ligation assay (PLA) was performed in pre-extracted cells using anti-PCNA antibody and RNAPII-pS2 as markers of active replication and transcription, respectively. Strikingly, EdU-positive CGGBP1-depleted cells showed a substantial increase in TRC-PLA foci compared to control knockdown cells (**Figures 4A-B**). We also observed a substantial decrease of EdU incorporation as a measure of DNA synthesis rates in siCGGBP1 versus siControl S-phase cells, suggesting that DNA replication is impaired in the absence of CGGBP1 (**Figure 4C**).

**Figure 4.**
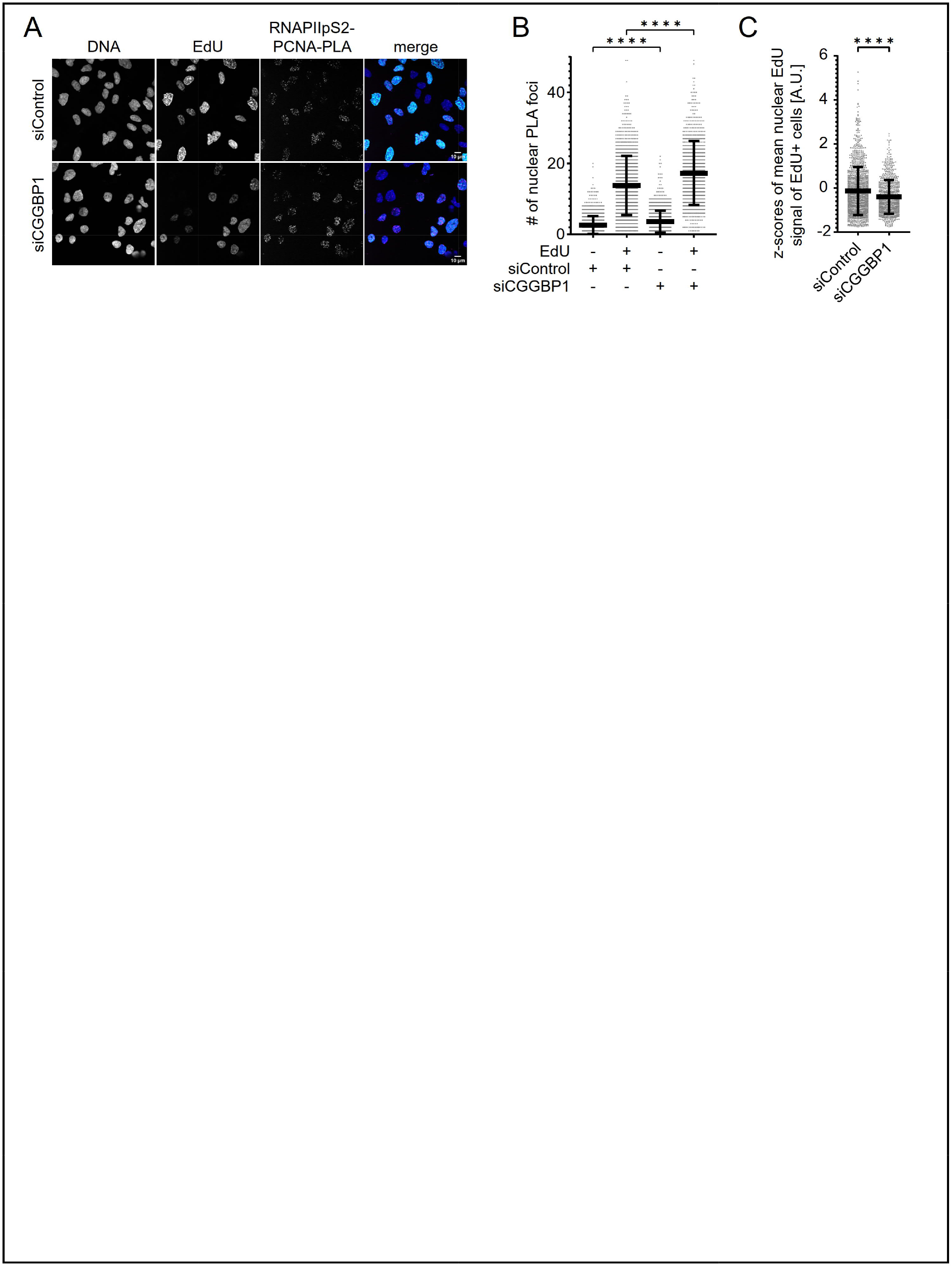
CGGBP1 levels are important to mitigate transcription-replication interference. A) Example IF images of EdU incorporation and RNAPII-pS2 – PCNA proximity ligation assay foci in U2OS cells upon 48 h CGGBP1 knockdown compared to control siRNA. N=2. B) Quantification of nuclear PLA foci in EdU- and EdU+ cells in all fields similar to A). Data are represented as mean ± standard deviation. Statistical significance was calculated using ordinary one-way ANOVA. N=2. C) Quantification of mean nuclear EdU signal of EdU+ cells in all fields similar to A). Data are represented as mean of the z-scores ± standard deviation. Statistical significance was calculated using two-tailed student’s t-test. N=2.

Given that CGGBP1 binds preferentially at actively transcribed promoter sites (**Figure 1**) and its depletion leads to elevated TRCs and replication stress (**Figure 4**), we investigated whether CGGBP1 co-localizes with initiating RNAPII using a specific antibody against RNAPII phospho-Ser5 (RNAPII-pS5). We observed significantly more RNAPII-pS5-CGGBP1 PLA foci per cell in comparison to the single antibody controls independently of the S-phase stage (**Figures S4A-B**), suggesting that CGGBP1 can directly interact with initiating RNAPII. Interestingly, we also observed an interaction of CGGBP1 with PCNA and RPA32-pS33 (**Figures S4C and S4C-D**). This result indicates that CGGBP1 may colocalize with sites of active replication and replication fork stalls in S-phase cells, further supporting a role of CGGBP1 at sites of transcription-induced replication interference.

### Pyridostatin-induced DNA damage in S-phase cells is dependent on cellular CGGBP1 levels

We next investigated the mechanism behind the observed transcription-induced replication stress in cells with altered CGGBP1 levels. We hypothesized that the binding of CGGBP1 at these short repeats may counteract or prevent the formation of secondary DNA structures such as G4 and R-loops. To test this hypothesis, we first treated cells with pyridostatin (PDS), a highly selective G-quadruplex–binding small molecule expected to stabilize intracellular G4 structures regardless of sequence variability (Müller et al., 2010; Rodriguez et al., 2008, 2012). It was previously reported that PDS can displace certain G4-binding proteins such as transcription factors due to the high binding affinity between PDS and G4s to disturb gene expression (Li et al., 2020, 2021).

However, the influence of PDS on CGGBP1 binding and genome-wide gene expression regulation have been unexplored. Therefore, we treated siControl and siCGGBP1-depleted cells with PDS and monitored nascent transcription with a short pulse of EU labelling and DNA damage by immunofluorescence against γH2AX. Interestingly, PDS-treated control knockdown cells showed a significant increase in EU incorporation (**Figures 5A-B**). Because G4s are bound by several transcription factors (Spiegel et al., 2021), PDS-binding to G4s may compete with the transcription factor landscape, thereby inducing the observed increase of EU incorporation. PDS treatment in siControl cells also led to increased γH2AX foci formation (**Figures 5A and 5C**), consistent with the model that PDS-stabilized G4s provoke double-strand breaks (DSBs) by forming a strong block to DNA or RNA polymerases (Belotserkovskii et al., 2017; Broxson et al., 2011; De and Michor, 2011; Lemmens et al., 2015; Sarkies et al., 2010, 2012). Unexpectedly, CGGBP1 depletion in PDS-treated cells largely rescued these effects, leading to less EU incorporation and a reduction of γH2AX foci formation compared to siControl cells (**Figures 5A-C**). This reduction of EU incorporation primarily occurred in non-nucleolar regions, indicating that CGGBP1 mainly affected nuclear transcription rates upon PDS treatment (**Figure S5A-B**). Together, this suggests that the presence of CGGBP1 at PDS-stabilized G4 structures may result in a stronger impediment to the replication and transcriptional machinery, leading to more DNA damage.

**Figure 5.**
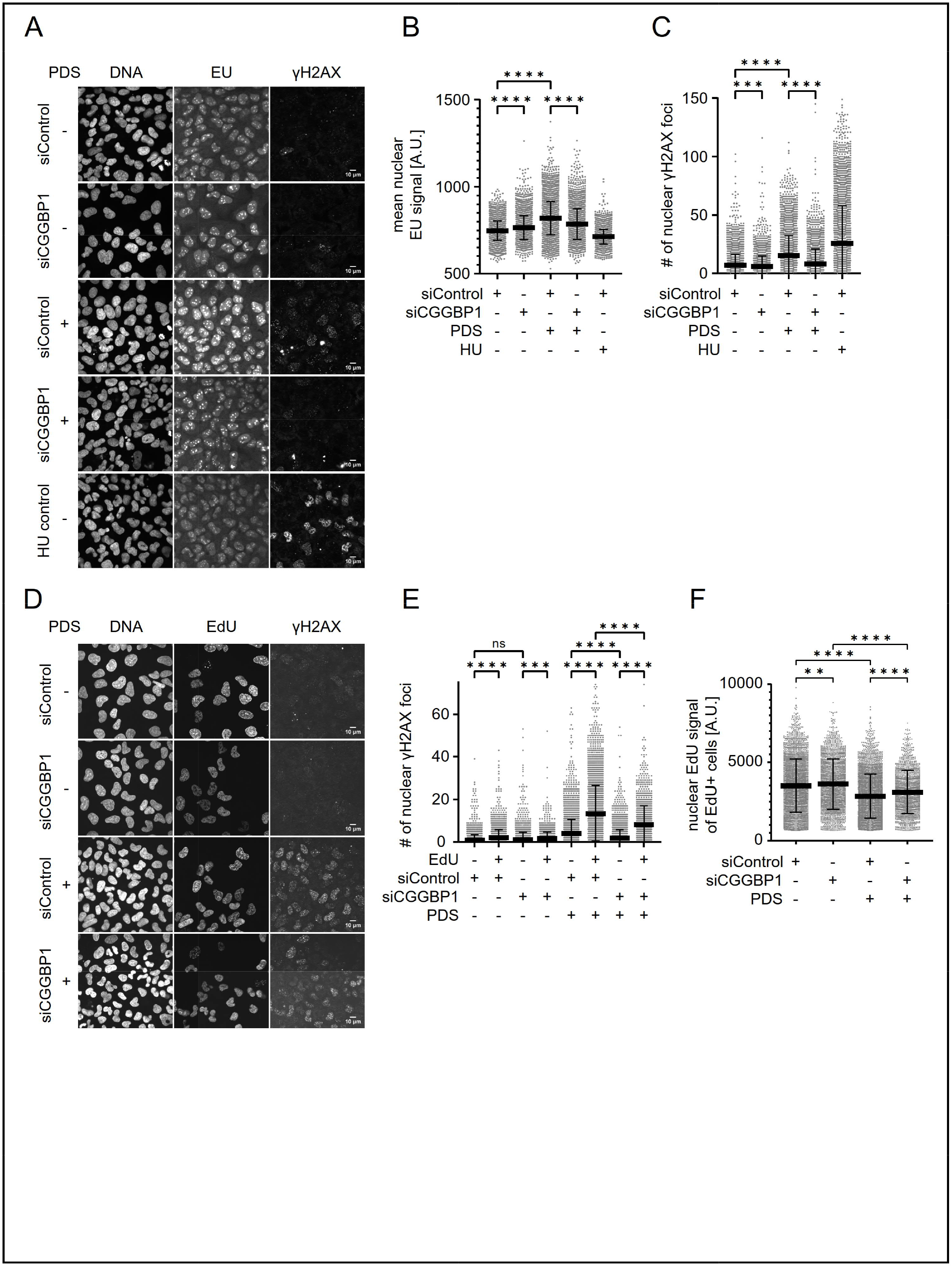
Pyridostatin-induced DNA damage in S-phase cells is dependent on cellular CGGBP1 levels. A) Example IF images of γH2AX signal and EU incorporation in U2OS cells upon 72 h CGGBP1 knockdown compared to control siRNA. Cells were treated with 20 μM pyridostatin (PDS) or with 500 μM hydroxyurea for 4h or 2h before fixation, respectively. N=1. B) Quantification of mean nuclear EU signal in all fields similar to A). The data of siControl and siCGGBP1 without treatment are already shown in Figure 2D, but plotted here again for comparison reasons. Data are represented as mean ± standard deviation. Statistical significance was calculated using ordinary one-way ANOVA. N=1. C) Quantification of nuclear γH2AX foci in all fields similar to A). Data are represented as mean ± standard deviation. Statistical significance was calculated using ordinary one-way ANOVA. N=1. D) Example IF images of γH2AX signal and EdU incorporation in U2OS cells upon 72 h CGGBP1 knockdown compared to control siRNA. Cells were treated with 20 μM PDS for 4h before fixation. N=1. E) Quantification of nuclear γH2AX foci in all fields similar to D). Cells of the same condition are split into EdU- and EdU+. Data are represented as mean ± standard deviation. Statistical significance was calculated using ordinary one-way ANOVA. N=1. Further uantification in **Figure S5A**. F) Quantification of mean nuclear EdU signal of EdU+ cells in all fields similar to D). Data are represented as mean ± standard deviation. Statistical significance was calculated using ordinary one-way ANOVA. N=1.

To further test the dependency of this effect on DNA replication, we repeated the experiment and included a short pulse of EdU in order to distinguish between cells in S-phase and non-S-phase cells. PDS-induced γH2AX foci were particularly high in EdU-positive siControl cells (**Figures 5D-E**), suggesting that a large fraction of PDS-induced DNA damage occurs during DNA replication. Consistently, we also observed less EdU-positive cells (**Figure S5C**) and a major reduction in EdU incorporation upon PDS treatment in S-phase cells (**Figure 5F**). Importantly, CGGBP1 depletion efficiently decreased γH2AX foci particularly in S-phase cells and could partially restore EdU incorporation (**Figure 5D-F**), supporting the notion that the combined binding of CGGBP1 at PDS-stabilized G4 structures is a major roadblock to replication forks and therefore causes DNA damage checkpoint activation and cell cycle arrest.

### R-loops are enriched at a CGG-repeat containing episomal transcription unit after CGGBP1 depletion

To probe the consequences of CGGBP1 binding on the formation of co-transcriptional R-loops, we took advantage of a previously established episomal system to induce R-loop dependent TRCs (Hamperl et al., 2017). In order to provide a suitable substrate for CGGBP1 binding at an active promoter site, we introduced 10 CGG repeats ((CGG)_10_) downstream of the doxycycline (DOX)-inducible Tet-ON promoter to allow for controlled transcription over the (CGG)_10_ repeat sequence. Because a similar approach with another R-loop forming sequence showed robust R-loop accumulation only in the HO orientation (Hamperl et al., 2017), we inserted the (CGG)_10_ sequence in a head-on (HO) orientation to the unidirectional Epstein–Barr virus replication origin oriP (**Figure 6A**). We then generated monoclonal human U-2OS cell lines with a stable integration of ∼5-70 plasmid copies per cell (**Figure S6A–B**).

**Figure 6.**
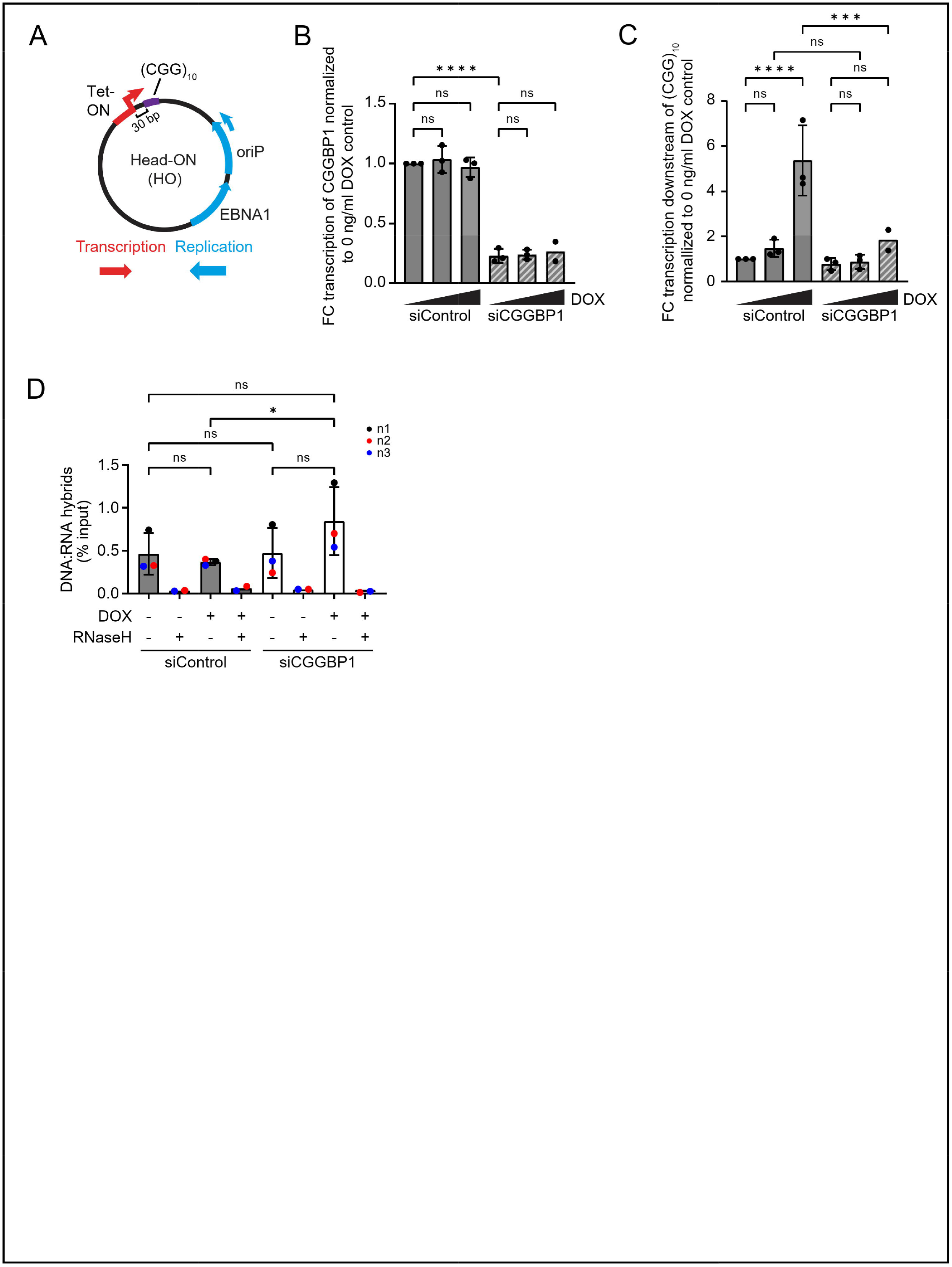
CGGBP1 binding opposes secondary structure formation and promotes transcription elongation on an episomal system. A) Plasmid map of the episomal transcription-replication conflict system with 10 CGG repeats inserted at the Tet-ON promoter. EBNA1 forces unidirectional replication at the oriP. Doxycycline treatment induces transcription leading to head-on conflicts with replication. B) Gene expression of CGGBP1 measured by RT-qPCR of cDNA from U2OS Tet-ON pHU43 clone 2 carrying the episomal system and treated with siControl or siCGGBP1 for 72 h. Cells were treated with 0, 100 or 1000 ng/ml DOX for 72 h. Shown is the fold change relative to MCM3 and normalized to the 0 ng/ml DOX control. Data are represented as mean ± standard deviation. Statistical significance was calculated using ordinary one-way ANOVA. N=3, except for the siCGGBP1 1000 ng/ml condition. See **Figure S6A-B** for characterization of U2OS Tet-ON pHU43 clone 2. C) Gene expression downstream of (CGG)_10_ measured by RT-qPCR of cDNA from U2OS Tet-ON pHU43 clone 2 carrying the episomal system and treated with siControl or siCGGBP1 for 72 h. Cells were treated with 0, 100 or 1000 ng/ml DOX for 72 h. Shown is the fold change relative to MCM3 and normalized to the 0 ng/ml DOX siControl condition. Data are represented as mean ± standard deviation. Statistical significance was calculated using ordinary one-way ANOVA. N=3, except for the siCGGBP1 1000 ng/ml condition (N=2). D) DNA:RNA hybrid immunoprecipitation (DRIP) with the S9.6 antibody downstream of the (CGG)_10_ repeat measured by qPCR of U2OS Tet-ON pHU43 clone 2 treated with siControl or siCGGBP1 and 0 or 1000 ng/ml DOX for 72h. Data are represented as mean ± standard deviation. Statistical significance was calculated using ordinary one-way ANOVA. N=3, except for RNaseH controls (N=2).

We performed siRNA-mediated knockdown of CGGBP1 for 72h (**Figure 6B**) and simultaneously induced transcription of (CGG)_10_. RT-qPCR indicated dose-dependent transcriptional activation of Tet-ON controlled (CGG)_10_ in control knockdown cells (**Figure 6C**). Interestingly, transcription through (CGG)_10_ was strongly reduced in CGGBP1-depleted cells compared to control cells (**Figure 6C**). This could indicate a role for CGGBP1 in efficient RNAPII translocation through the plasmid-derived CGG repeats. Importantly, this transcriptional defect could not be explained by changes in the plasmid copy number per cell as this remained unchanged at the different DOX concentrations (**Figure S6C**). To determine if this transcriptional defect caused upon CGGBP1 depletion stems from R-loop formation on the (CGG)_10_ sequence in cells, we performed DNA-RNA immunoprecipitation (DRIP) and qPCR on the plasmid using the RNA-DNA hybrid-specific S9.6 antibody (Boguslawski et al., 1986). The siControl cells exhibited a certain baseline level of RNase H-sensitive RNA-DNA hybrid formation, which was not significantly increased upon DOX induction (**Figure 6D**), in line with previous data showing that non-disease short tracts of CGG repeats at the FMR1 locus do not show efficient R-loop formation upon their transcriptional activation (Groh et al., 2014). Importantly, RNA:DNA hybrids were significantly induced in the CGGBP1-depleted cells after DOX induction (**Figure 6D**). However, the transcriptional activity of the reporter gene remained reduced under (**Figure 6C**). This finding suggests that the binding of CGGBP1 to short CGG repeats counteracts or prevents the formation of RNA:DNA hybrids while these repeats undergo transcription, thereby enabling efficient mRNA production.

Together, these data provide clear evidence that CGGBP1 is a critical factor that controls the formation of genome-destabilizing R-loop structures that can form over short CGG repeats enriched at gene promoter sites. In the absence of CGGBP1, the resulting DNA secondary structures may form a structural block, directly interfering with both RNAPII transcription elongation and replication fork progression.

## Discussion

### Short CGG tandem repeats prone to G4 and R-loop formation are enriched at gene promoter sites bound by CGGBP1

Short (CGG)n tandem repeats with n < 10 are frequent DNA motifs commonly found throughout the genome of higher eukaryotes (Willems et al, 2014) without any major impact on DNA replication fork progression or transcription elongation. However, i*n vitro* studies suggest that oligonucleotides with only 5-7 CGG repeats are able to form stable G4 structures (Fry and Loeb, 1994), which were demonstrated to be efficient replication impediments *in vivo* (Sarkies et al., 2012). In addition, CGG trinucleotide repeats have an inherent sequence bias towards high GC content and GC-skew, two features well known to promote the formation of co-transcriptional R-loop structures (Ginno et al., 2012, 2013). This raises an important conceptual question whether this apparently inert behaviour of CGG repeats *in vivo* is due to these sequences being incapable of forming secondary structures *in vivo* or whether it is the result of other interacting proteins that counteract structure formation at these repeat sequences. We now show in the context of CGG repeats that these sequences are enriched in the human genome at gene promoter sites that are prone to G4 and R-loop secondary structure formation and identify the CGG-sequence specific DNA-binding factor CGGBP1 as a specific interactor of these sequences *in vivo* (**Figure 1A-C**). It is important to note that despite clear enrichment of a CGG motif (**Figure 1D**), not all identified CGGBP1 ChIP-Seq peaks were directly associated with the presence of short CGG repeats, suggesting that CGGBP1 binding to chromatin may also be affected by additional chromatin features such as an interaction with nucleosomes or other chromatin components. Consistently, earlier studies identified CGGBP1 as part of a heterotrimeric complex with NFIX and the high-mobility group protein HMGN1 that acts as a bidirectional regulator of HSF1 transcription (Singh et al., 2009). Thus, specific binding partners of CGGBP1 could mediate additional interactions with certain chromatin features and therefore expand the genomic CGGBP1 target sites. Nevertheless, the enrichment of CGGBP1 at gene promoters was clearly observed irrespective of its precise targeting or binding mechanism.

### Altering cellular CGGBP1 levels leads to transcription elongation impairment and replication fork stalling

Previous work has demonstrated that increased levels of DNA secondary structures such as G4 and co-transcriptional RNA:DNA hybrids provide roadblocks for both RNA and DNA polymerases, resulting in transcription elongation impairment and replication fork slow-down and stalling (Belotserkovskii and Hanawalt, 2015; Gan et al., 2011; Groh et al., 2014; Tous and Aguilera, 2007). Consistently, we observed that CGGBP1-depleted cells show a transcription- and S-phase-dependent accumulation of active RNAPII on chromatin (**Figure 3**), which resulted in replication fork impediment and increased transcription-replication interference as measured by a TRC-PLA assay (**Figure 4**). Notably, we observed an increase in nascent transcription as measured by EU incorporation in CGGBP1-depleted cells (**Figure 2A-B**), which is difficult to reconcile with our model of RNAPII transcription elongation impairment in S-phase. One potential explanation is that CGGBP1 additionally regulates genes transcribed by other RNA polymerases. In fact, ribosomal RNA (rRNA) genes transcribed by RNAPI are rich in CGG repeats (Müller-Hartmann et al., 2000) and CGGBP1 shows preferential binding to rRNA gene clusters on the small arms of acrocentric chromosomes, which is consistent with our finding that EU signal is increased in nucleoli after CGGBP1 knockdown (**Figure S5A-B**). In addition, CGGBP1 was found to bind RNAPIII promoters, acting as a transcription inhibitor particularly at the repetitive Alu-SINE elements (Agarwal et al., 2014). Together, the observed global increase of EU incorporation in CGGBP1-depleted cells may partially be explained by the changed transcription dynamics of these two other nuclear RNA polymerases. Conversely, CGGBP1 overexpression led to a global decrease in EU incorporation as well as a decrease of chromatin-bound RNAPII (**Figure 2** **and** **Figure 3**), suggesting that excess CGGBP1 levels has an inhibitory effect most likely on all nuclear RNA polymerases. Together, these data suggest that a balanced level of cellular CGGBP1 levels is critical for efficient nuclear transcription rates, but the detailed mechanism how CGGBP1 controls transcription elongation rates of the different RNA polymerases is likely complex and may also be caused by secondary effects due to the cell cycle arrest or DNA damage checkpoint activation in CGGBP1-depleted cells.

### Promoter-bound CGGBP1 slows RNAPII elongation and prevents G4/R-loop formation

Instead, the focus of this study was to analyze the role of CGGBP1 in preventing or counteracting DNA secondary structure formation. For this, we used two distinct approaches in order to test the role of CGGBP1 in G4 or R-loop resolution. Stabilization of G4 structures by PDS caused replication impairment and DNA damage in S-phase cells, which could be partially rescued by CGGBP1 depletion (**Figure 5**). This surprising finding suggests that the combined presence of CGGBP1 at PDS-stabilized G4 structures may result in a stronger roadblock to replication forks and therefore cause more replication stress and DNA damage and provides a clear connection between CGGBP1 levels and G4-induced DNA damage. Interestingly, CGGBP1 has a C-terminal SQ motif that was identified as a substrate of the ATR DNA damage response kinase (Matsuoka et al., 2007; Singh and Westermark, 2015; Traven and Heierhorst, 2005). Thus, CGGBP1 may have a functional role in DNA checkpoint signalling of G4-induced DNA damage, but more detailed studies will be necessary to understand the mechanistic details of this interaction.

Finally, we took advantage of an episomal plasmid system with a short transcription-inducible CGG repeat and show that CGGBP1 depletion blocks transcription and induces the formation of RNA:DNA hybrid structures on the repeat sequence (**Figure 6**), providing strong evidence that CGGBP1 counteracts or prevents the formation of RNA:DNA hybrids while these repeats undergo transcription, thereby inhibiting efficient mRNA production. Together, we propose a model that short trinucleotide repeats can give rise to genome-destabilizing secondary structures and rely on specific cellular factors such as CGGBP1 to maintain accurate transcription and replication programs (**Figure 7**). In the future, it will be interesting to understand if other trinucleotide repeat sequences such as GAA repeats that show similar replication fork impediments (Šviković et al., 2019) and human disease-relevant instabilities (Den Dunnen, 2018; Jones et al., 2017) also rely on the interaction of sequence-specific binding partners to reduce secondary structure formation, thereby playing important roles in transcription of the repeats and preventing clashes between transcription and replication.

**Figure 7.**
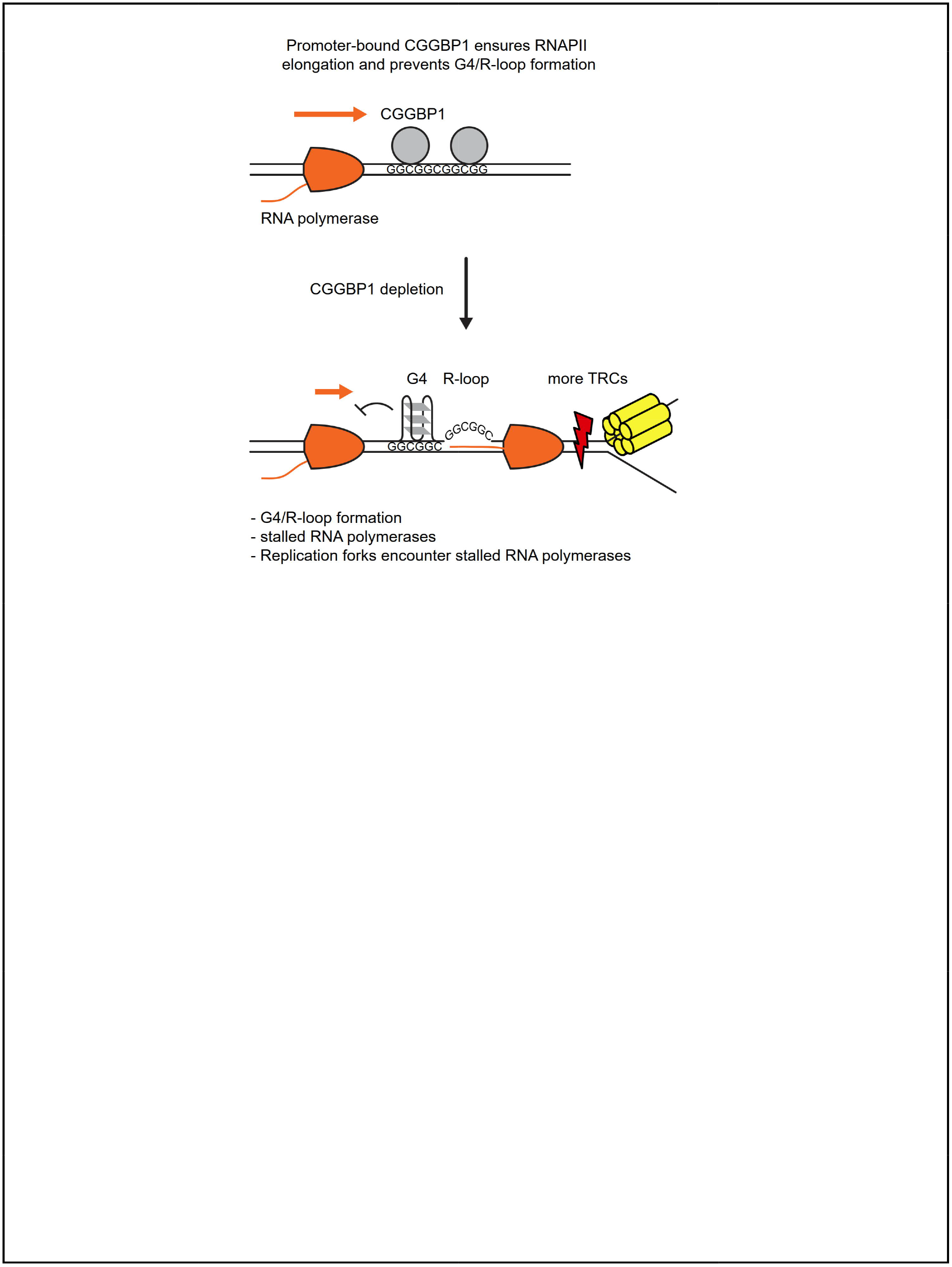
Model of how CGGBP1 binding at short CGG repeats opposes secondary structure formation and promotes transcription elongation. The presence of CGGBP1 at promoter sites prevents the formation of DNA secondary structures capable of blocking RNA polymerases and replication forks. Depletion of CGGBP1 leads to the formation of DNA secondary structures resulting in the accumulation of stalled RNA polymerases. In S-phase, replication forks encounter the stalled RNA polymerases, leading to unscheduled transcription-replication conflicts and replication impairment.

## Tables with titles and legends

**Supplementary Table 1.**
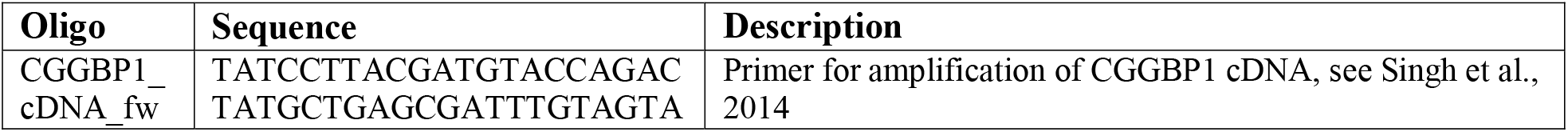
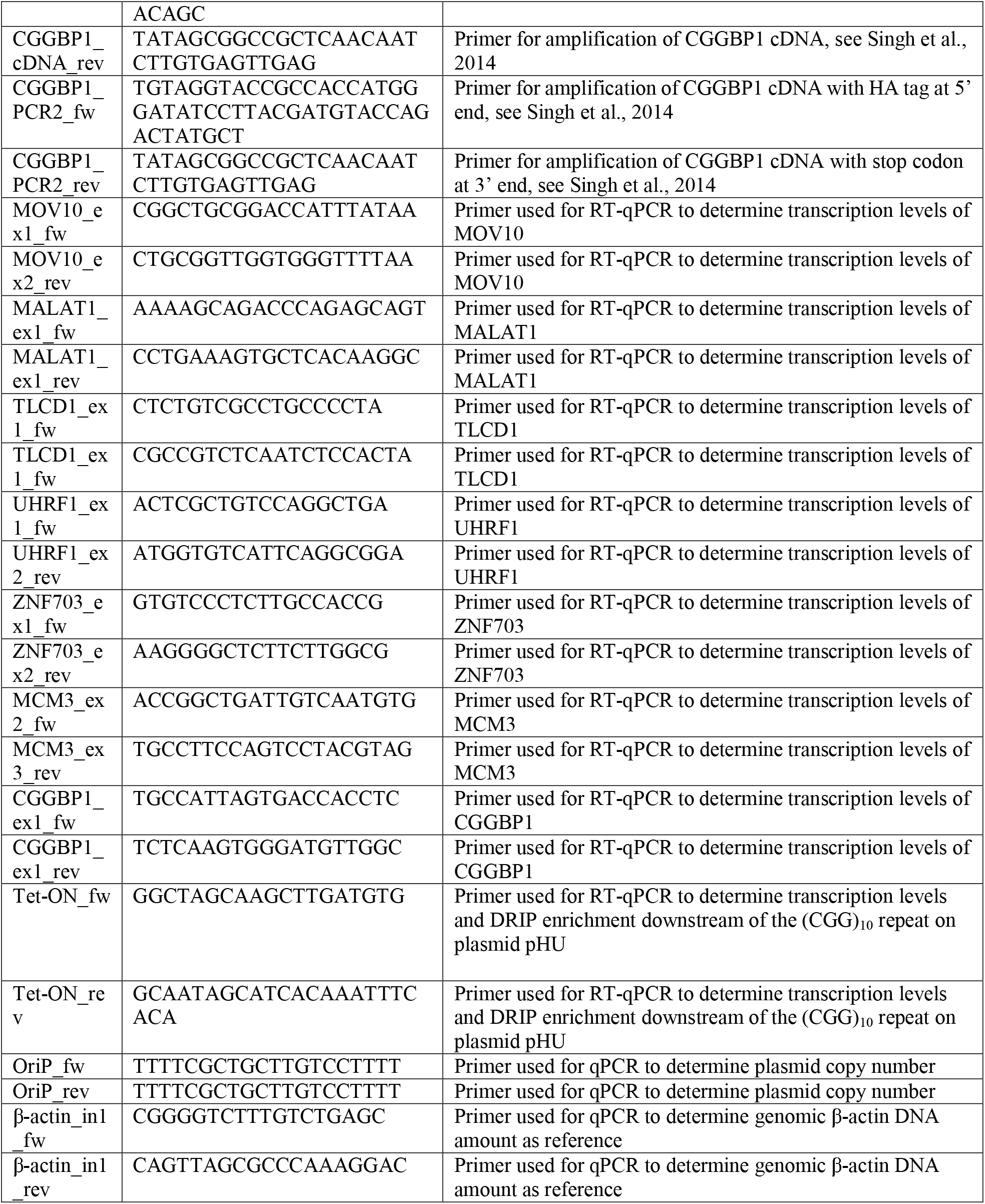
Oligonucleotides used in this study.

**Supplementary Table 2.** Plasmids used in this study.

| Plasmid | Insert | Construction/Source |
| --- | --- | --- |
| pcDNA3.1+ | - | Thermo Fisher Scientific (Cat. #V79020) |
| pcDNA3.1 HA-CGGBP1 | HA-CGGBP1 | Cloned as described in Singh et al., 2014 |
| pHU43 | (CGG)<sub>10</sub> | Insertion of KpnI and HindIII cut Q-block fragment containing (CGG)<sub>10</sub> into pSH36 from Hamperl et al., 2017. Cloned following Scior et al., 2011 |

## Acknowledgements

We thank the ENCODE production laboratory for the CGGBP1 ChIP-Seq dataset, Adam Burton and Maria-Elena Torres-Padilla for discussions and comments on the manuscript. S.H. is supported by the European Research Council (ERC Starting Grant 852798 ConflictResolution) and by the Deutsche Forschungsgemeinschaft (CRC1064).

## Author Contributions

Conceptualization and methodology, H.U., E.K. and S.H.; formal analysis, H.U. and S.H.; data curation and software, M.T., M.L., A.E; visualization, H.U., and S.H.; funding acquisition, project administration, supervision, and resources, S.H. investigation and validation, H.U., M.L., M.T., M.W., E.K., A.C., M.W., C.L. and S.H. writing (original draft), S.H.; writing (review and editing), H.U., M.L., M.T., M.W., A.C, C.L., E.K., M.W., and S.H.

## STAR Methods

### Cell lines

Human osteosarcoma-derived cells (U2OS) that express the tetracycline regulated transactivator Tet-ON were used for all experiments. U2OS Tet-ON cells were cultured in DMEM (GIBCO) supplemented with 10% Tet-ON approved FBS, 4.5 g/L D-Glucose and L-Glutamine, 110 mg/L sodium pyruvate and penicillin/streptomycin in 5% CO2 at 37°C. To generate monoclonal U2OS Tet-ON cell lines, cells were transfected with the vector pHU43 ((CGG)_10_ -HO) and selected with 200 µg/ml hygromycin B for 2-3 weeks. Surviving cells were harvested and single cells were sorted into 96-wells by flow cytometry (BD FACSMelody) and cultured to expand over 5-6 weeks under selection in 200 µg/ml hygromycin.

### Oligonucleotides and plasmids

Standard techniques were used for cloning of plasmids. Complete lists of oligonucleotides and plasmids used in this study can be found in Supplementary Tables 1 and 2.

### Plasmid and siRNA transfections

TransIT-LT1 (Mirus) or Lipofectamine RNAiMAX (ThermoFisher) transfection reagents were used for all plasmid and siRNA transfections, respectively, following the manufacturer’s instructions. For plasmid transfections, a ratio of 2 µg plasmid DNA per 6 µL transfection reagent was used and the mixture was incubated for 30min before adding dropwise to cells. For siRNA transfections, a ratio of 50 nM siRNA per 2 µL transfection reagent was used and the mixture was incubated for 20min before adding dropwise to cells. For all transfections, culture media was replaced with fresh media after 12-16 h incubation at 37°C.

### Chromatin fractionation and Western blotting

Chromatin fractionation was performed as described (Méndez and Stillman, 2000) with minor changes. In brief, 1 x 106 cells were resuspended in buffer A (10 mM HEPES pH7.9, 10 mM KCL, 1.5 mM MgCl2, 340 mM sucrose, 10% Glycerol, 1 mM DTT) and lysed by incubating with Triton X-100 at a final concentration of 0.1% for min on ice. Nuclei were collected by low-speed centrifugation at 1300 g for 4min and 4°C and washed once in buffer A before lysing in buffer B (3 mM EDTA, 0.2 mM EGTA, 1 mM DTT, 1x protease inhibitor cocktail) for 30min on ice. Insoluble chromatin was collected by centrifugation at 1700 g for 4 min and 4 °C and washed once with buffer B before resuspending final chromatin pellet in 2x LDS sample buffer.

For whole cell extracts, cells were lysed in RIPA buffer (50mM Tris HCl, pH 7.4, 150mM NaCl, 0.5% deoxycholate, 0.1% sodium dodecyl sulfate, 1% NP-40, 1x protease inhibitor cocktail). Whole cell extracts and chromatin fractions were sonicated alternating 2.5s on, 2.5s off for 10 min (Bioruptor UCD-200) before separation by eletrophoresis and transfer onto polyvinylidene difluoride membranes. Membranes were blocked in 5% skimmed milk dissolved in 0.1% Tween/PBS overnight at 4°C. Membranes were incubated with primary antibodies dissolved in 3% BSA/PBS overnight at 4°C followed by washing thrice with 0.1% Tween/PBS. HRP-linked secondary antibodies were then added for 1h at 25°C and washed thrice prior to signal detection using ECL reagent. Primary antibodies:antibodies: CGGBP1 (1:1000), CDC45 (1:1000), GAPDH (1:10000), total RNAPII (1:5000), RNAPII pS2 (1:5000), H3 (1:10000), γH2AX (1:300), pRPA32 S33 (1:2500).

### Immunofluorescence imaging

For all IF experiments, except for EU incorporation and γH2AX staining, cells were pre-extracted with CSK100 buffer (100 mM NaCl, 300 mM Sucrose, 3 mM MgCl2, 10 mM MOPS, 0.5% Triton X-100 in PBS) and washed once with PBS. Cells were fixed with 4% PFA/PBS for 15 min, washed once with PBS, permeabilized with 0.5% Triton X-100 for 10 min and washed once more. For EdU or EU staining, cells were incubated with 10 μM EdU or 0.5 mM EU 30 min before fixation and a commercial kit was used to attach fluorescent dyes with a click reaction (ThermoFisher). Cells were blocked in 3% BSA/PBS for 1h at 25°C. Primary antibodies diluted in 3% BSA/PBS were added overnight followed by three PBS washes. Secondary antibodies diluted in 3 % BSA/PBS were added together with DAPI (5 μg/ml) for 90min followed by three PBS washes. Cells were imaged at 40x on a spinning disc confocal microscope (Andor Dragonfly). Images were analyzed using a custom ImageJ script that allows nuclei segmentation on the DAPI channel with StarDist (Schmidt et al., 2018). To create masks of the nucleoli in the EU channel, threshhold_yen from the scikit-image python package was used (van der Walt et al., 2014; Yen et al., 1995). Primary antibodies: total RNAPII (1:2000), RNAPII pS2 (1:2000), HA-tag (1:1000), γH2AX (1:300). Secondary antibodies: anti-mouse-Alexa 594 (1:1000) Anti-rabbit-Alexa 647 (1.1000).

### Cell cycle analysis

Cells were pulse-labeled with 25 μM 5-Bromo-2′-deoxyuridine (BrdU) for 30 min, and washed once with PBS before trypsinization and harvesting. After fixing samples with ice-cold 70% ethanol, cells were permeabilized with 0.25% Triton X-100/PBS for 15 min on ice. To denature DNA to single strand, cells were incubated in 2N HCl for 15min at 25°C followed by a wash with 100 mM sodium borate pH 8.5. Cells were blocked in 1% BSA/PBS containing 0.1% Tween-20 for 15 min, and incubated in primary BrdU antibody (1:100) for 2 h. Cells were then washed three times in PBS, incubated in AlexaFluoro-488 secondary antibody for 1 h, and washed three times with PBS. Propidium iodide (PI; 0.1 mg/mL; Sigma) and RNase A (10 mg/mL) was added to determine DNA content and cells were analyzed on a FACSMelody device (BD Bioscience). Cell cycle profiles were determined using FlowJo software.

### Reverse transcription-qPCR

Cells were harvested and total RNA was isolated using TRIzol reagent following the manufacturer’s protocol. After digestion with RNase-free DNase I at 37°C for 30min, reverse transcription was carried out with 0.5-1.0 μg total RNA with random hexamer primers and SuperScript III Reverse Transcriptase Kit. Equal amounts of cDNA were mixed with iTaq SYBR Green Supermix and run on a Roche LightCycler 480 Instrument II. mRNA expression levels were measured by the change in comparative threshold cycles with primer pairs in Table S1 and MCM3 cDNA as a reference.

### Proximity ligation assay (PLA)

Cells were pre-extracted with CSK100 buffer (100 mM NaCl, 300 mM Sucrose, 3 mM MgCl2, 10 mM MOPS, 0.5% Triton X-100 in PBS) and washed once with PBS. Cells were fixed with 4% PFA/PBS for 15 min, washed once with PBS, permeabilized with 0.5% Triton X-100 for 10 min and washed once more. For EdU staining, a commercial kit was used to attach fluorescent dyes with a click reaction (ThermoFisher). Cells were blocked in 5% BSA/PBS for 1h at 25°C and incubated with primary antibodies diluted in 5 % BSA/PBS overnight at 4°C followed by two PBS washes. The PLA was performed following the manufacturer’s protocol (Sigma) with slight changes. In Brief, cells were incubated for 1h at 37°C with 1:10 diluted PLA probes in Duolink antibody diluent followed by two 5min washes in wash buffer A. To ligate the probes, ligase was added 1:70 to 1x ligation buffer and cells were incubated for 30min at 37°C followed by two 5 min washes in wash buffer A. For amplification, polymerase was added 1:140 to 1x amplification buffer and cells are incubated for 100min at 37°C followed by two 10 min washes in wash buffer B. For DNA staining, DAPI (5 µg/ml) diluted in PBS was added for 90 min before washing twice with PBS. Cells were imaged at 40x on a spinning disc confocal microscope (Andor Dragonfly). The number of PLA foci was quantified using ImageJ after nuclei segmentation on the DAPI channel. Primary antibodies: RNAPII pS2 (1:2000), PCNA (1:2000), RNAPII pS5 (1:2000), CGGBP1 (1:100), pRPA32 S33 (1:2000).

### Plasmid copy number analysis

Genomic DNA was isolated from 2-4 × 105 cells. After trypsinization, cells were washed in 1x PBS and resuspended in TE buffer followed by the addition of an equal volume of IRN buffer (50mM Tris-HCl at pH 8, 20 mM EDTA, 0.5 M NaCl), 0.5% SDS and 10 μg Proteinase K. After digestion for 1h at 37°C, DNA was extracted with phenol/chloroform and digested with 20 μg RNase A for 1h at 37°C. After chloroform extraction, DNA was precipitated with EtOH/sodium acetate, washed with 70% EtOH, and resuspended in TE. DNA was digested with EcoRI (NEB) restriction enzyme overnight at 37°C. The plasmid copy number was analyzed with primer pairs in Table S1 by amplifying either the oriP region of the plasmid or a region of the genomic beta-actin gene. The relative plasmid copy number was determined by quantitative PCR on a Roche LightCycler 480 Instrument II using SYBR-Green master mix (Biorad) and defined as the ratio of the amount of 2x oriP to the amount of beta-Actin.

### DNA:RNA immunoprecipitation

DRIP was performed as described in Ginno et al., 2012 with minor changes. Briefly, DNA was extracted with phenol/chloroform, precipitated with EtOH/sodium acetate, washed with 70% EtOH, and resuspended in TE. DNA was sonicated with 10% duty factor, 200 cycles/burst, 140 peak incident power for 4 min (Covaris). For RNase H-treated samples, 4 ug of DNA was treated with RNase H overnight at 37°C. DNA was purified by phenol/chloroform, EtOH/sodium acetate precipitation as described above. 4 μg of DNA was bound with 7 μg of S9.6 antibody in 1 X binding buffer (10 mM NaPO4 pH 7, 140 mM NaCl, 0.05% Triton X-100) overnight at 4°C. Protein G magnetic beads were added for 2 h. Bound beads were washed 3 times in binding buffer and elution was performed in elution buffer (50 mM Tris pH 8, 10 mM EDTA, 0.5% SDS, Proteinase K) for 45 min at 55°C. DNA was purified as described. Quantitative PCR of immunoprecipitated DNA fragments was performed on a Roche LightCycler 480 Instrument II using SYBR-Green master mix (Biorad).

### ChIP-seq data analysis

For CGGBP1 ChIP-seq analysis, we used publicly available ENCODE data of K562 cells with the following identifiers: ENCSR763FNU and ENCSR334KTB (Luo et al., 2020; The ENCODE Project Consortium, 2012). For most analysis, tools from the web platform galaxy were used (Afgan et al., 2018, https://usegalaxy.eu). To annotate CGGBP1 peaks, bed files were analyzed by ChIPseeker. For metagene plots of CGGBP1 ChIP-seq signal, bam files of the two replicates and the control were converted to bigWig with bamCoverage, heatmap values were calculated by deeptools computeMatrix and the plot created by deeptools plotHeatmap. For genome browser snapshots, these bigWig files were loaded into IGV (Robinson et al., 2011). For motif enrichment analysis, we used MEME-ChIP (Bailey et al., 2015). To analyze G4 hunter score, DRIP signal, GC% and GC skew a set of ∼2300 transcription start sites (TSS) that are proximal (<3kb away from promoter) to at least one CGGBP1 peak was selected. As controls, two sets of ∼2300 randomly selected TSS were created. To obtain G4 quadruplex formation prediction scores, the G4 Hunter python script was used on FASTA files of 2kb windows around the TSS with –w 25 –s 0.0 parameters (https://github.com/AnimaTardeb/G4-hunter.git). This calculates a mean G4 Hunter score at the middle of each 25 bp window sliding by 1bp across the 2kb. DRIP signal was obtained from a published data set (Castillo-Guzman et al., 2020, GSM4478670). The K562 DRIP-seq signal bigWig file was converted from build hg19 to hg38 with the CrossMap tool (Zhao et al., 2014). Heatmap values and the plot for the TSS sets were created by computeMatrix and plotHeatmap, respectively. For GC% and GC skew, a 10 kb window around the TSS was split into 200 bp windows using a sloding window of 1bp with bedtools MakeWindowsBed. Nucleotide content was calculated with bedtools NucBed and GC skew was then calculated by measuring (G-C)/(G+C).

### Quantification and statistical analyses

Statistical parameters including the number of biological replicates (N), standard deviation and statistical significance are reported in the figures and the figure legends. Statistical significance is determined by the value of p < 0.05 by One-Way ANOVA test, where appropriate. Where appropriate, we confirmed that sample sizes were large enough that any deviations from normality did not affect the statistical test results.

## Supplemental Information titles and legends

**Figure S1. Global profiling of CGGBP1 binding sites in the human genome**

A) Summary plot of GC% of the DNA sequence at either TSSs with a CGGBP1 binding peak in proximity or two random sets of TSSs.

B) Summary plot of GC skew of the DNA sequence at either TSSs with a CGGBP1 binding peak in proximity or two random sets of TSSs. GC skew was calculated with the formula (G-C)/(G+C).

**Figure S2. Altering cellular CGGBP1 levels leads to global transcriptional changes**

A) BrdU cell cycle analysis flow cytometry profile plots of U2OS cells upon treatment with siControl or siCGGBP1 for 24, 48 and 72 h. The percentage of cells in G1, S or G2 are provided next to the gates. N=1.

**Figure S3. Altering cellular CGGBP1 levels impacts the level of chromatin-bound RNAPII complexes**

A) Western Blot of U2OS chromatin fraction after treatment with siControl or siCGGBP1 for 72h. For transcriptional inhibition, 100 µM DRB or 1 µM triptolide were added 2 h before harvest. Depicted are total RNA polymerase II (RNAPII) and RNAPII pS2 detected on the same blot.

B) Quantification of mean integrated signal density relative to the H3 loading control in A). N=1.

C) Quantification of mean nuclear total RNAPII signal in all fields similar to **Figure 3A**. Data are represented as mean ± standard deviation. Statistical significance was calculated using ordinary one-way ANOVA. N=1.

D) Quantification of mean nuclear RNAPII-pS2 signal in all fields similar to **Figure 3A**. Data are represented as mean ± standard deviation. Statistical significance was calculated using ordinary one-way ANOVA. N=1.

E) Quantification of mean nuclear total RNAPII signal in all fields similar to **Figure 3A**. Cells of each condition are split into EdU- and EdU+. Data are represented as mean ± standard deviation. Statistical significance was calculated using ordinary one-way ANOVA. N=1.

F) Quantification of mean nuclear RNAPII-pS2 signal signal in all fields similar to **Figure 3A**. Cells of each condition are split into EdU- and EdU+. Data are represented as mean ± standard deviation. Statistical significance was calculated using ordinary one-way ANOVA. N=1.

G) Quantification of mean nuclear total RNAPII signal in all fields similar to **Figure 3D**. Cells of each condition are split into EdU- and EdU. Data are represented as mean ± standard deviation. Statistical significance was calculated using ordinary one-way ANOVA. N=1.

H) Quantification of mean nuclear RNAPII-pS2 signal in all fields similar to **Figure 3D**. Cells of each condition are split into EdU- and EdU. Data are represented as mean ± standard deviation. Statistical significance was calculated using ordinary one-way ANOVA. N=1.

**Figure S4. CGGBP1 levels are important to mitigate transcription-replication interference**

A) Example IF images of EdU incorporation and proximity ligation assay foci of the indicated antibody combination or single antibody controls in untreated U2OS cells. N=1.

B) Quantification of RNAPII-pS5 – CGGBP1 PLA foci in all fields similar to A). The single antibody controls are shown for both antibodies. Data are represented as mean ± standard deviation. Statistical significance was calculated using ordinary one-way ANOVA. N=1.

C) Quantification of PCNA – CGGBP1 PLA foci in all fields similar to A). The single antibody controls are shown for both antibodies. For comparison reasons, the same data of the single CGGBP1 antibody PLA control as in B) is shown. Data are represented as mean ± standard deviation. Statistical significance was calculated using ordinary one-way ANOVA. N=1.

D) Quantification of pRPA32(S33) – CGGBP1 PLA foci in all fields similar to A). The single antibody controls are shown for both antibodies. For comparison reasons, the same data of the single CGGBP1 antibody PLA control as in B) is shown. Data are represented as mean ± standard deviation. Statistical significance was calculated using ordinary one-way ANOVA. N=1.

**Figure S5. Pyridostatin-induced DNA damage in S-phase cells is dependent on cellular CGGBP1 levels**

A) Quantification of mean nucleolar EU signal in all fields similar to **Figure 5A**. Data are represented as mean ± standard deviation. Statistical significance was calculated using ordinary one-way ANOVA. N=1.

B) Quantification of mean nuclear EU signal excluding nucleoli in all fields similar to **Figure 5A**. Data are represented as mean ± standard deviation. Statistical significance was calculated using ordinary one-way ANOVA. N=1.

C) Bar graph showing the percentage of EdU+ and EdU-in all fields similar to **Figure 5D**. N=1

**Figure S6. CGGBP1 binding opposes secondary structure formation and promotes transcription elongation on an episomal system**

A) Initial plasmid copy numbers of different U2OS Tet-ON monoclonal cell lines carrying the episomal system measured by qPCR (see **Figure 6A**). Plasmid copy numbers were calculated with the relative ratio 2xOriP/β-actin. Data are represented as mean ± standard deviation. N=1.

B) Plasmid copy numbers of U2OS Tet-ON pHU43 clone 2 during passaging measured by qPCR. The time between passages was 3-4 days. DOX and siRNA treatment experiments were done between passages 1 and 4. Data are represented as mean ± standard deviation. N=1.

C) Plasmid copy number changes during DOX treatment of U2OS Tet-ON pHU43 clone 2. Data are represented as mean ± standard deviation. N=3.

