## Supplementary figures and images for "The CGG triplet repeat binding protein 1 counteracts DNA secondary structure-induced transcription-replication conflicts"

### Supplemental Figures

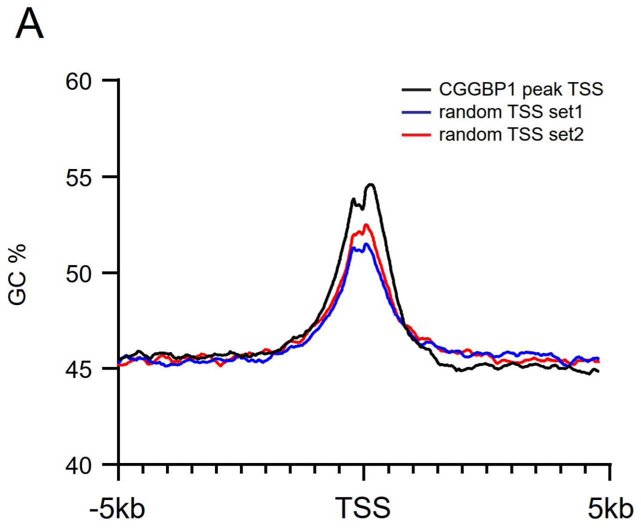

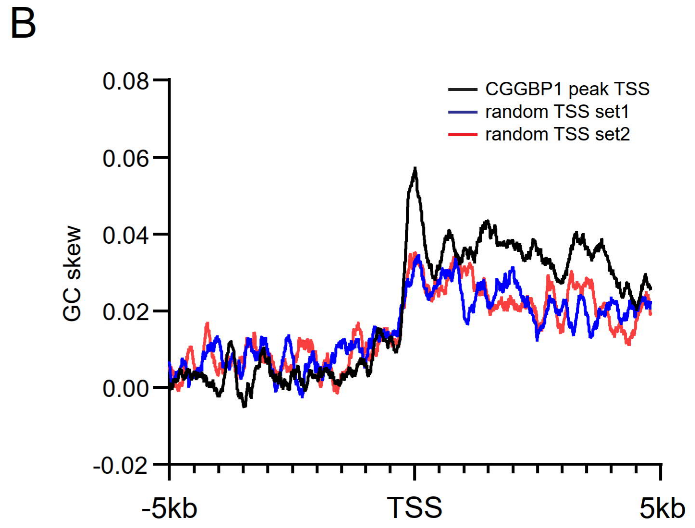

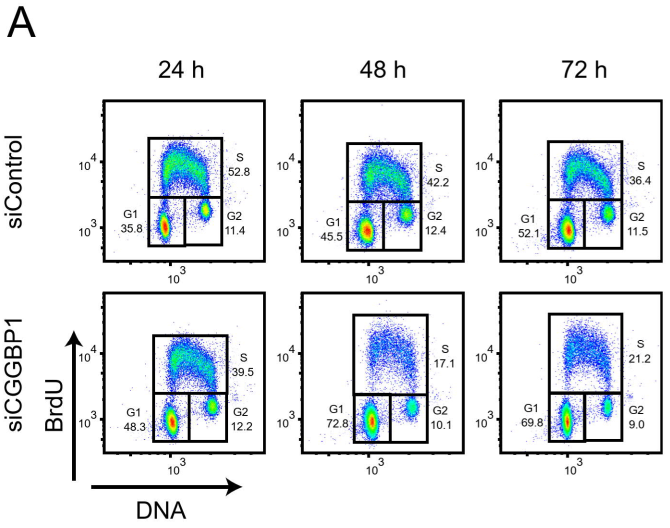

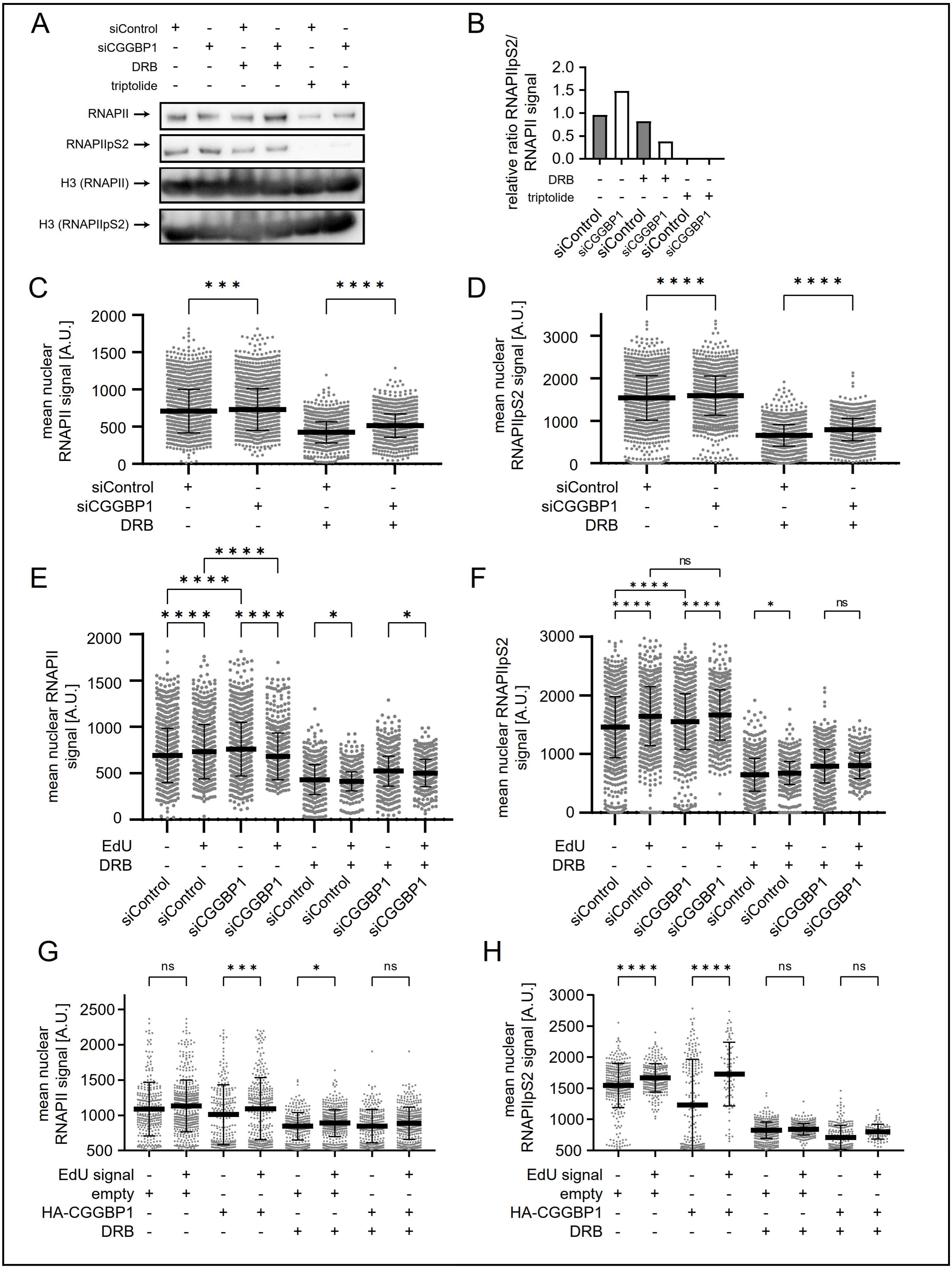

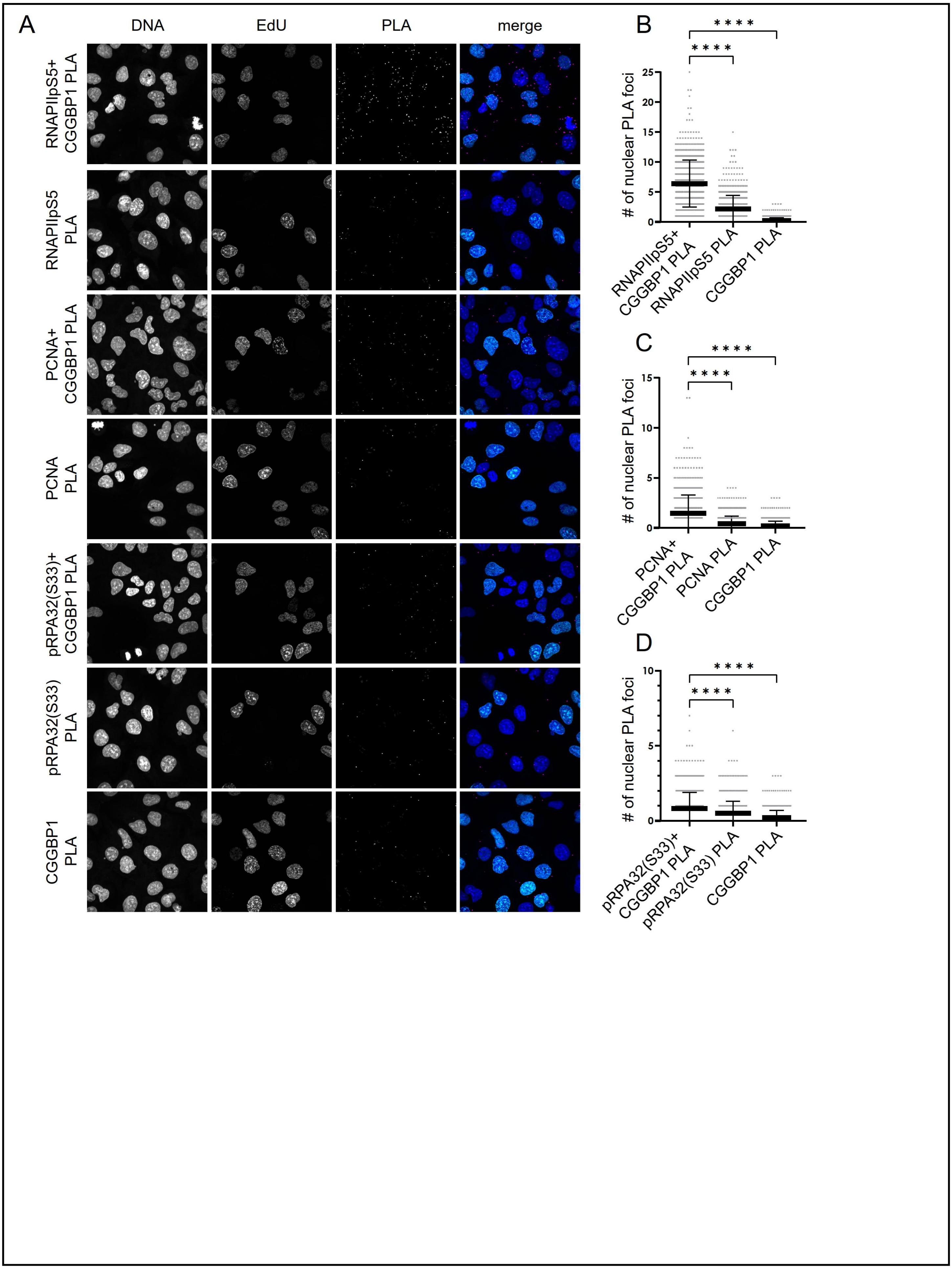

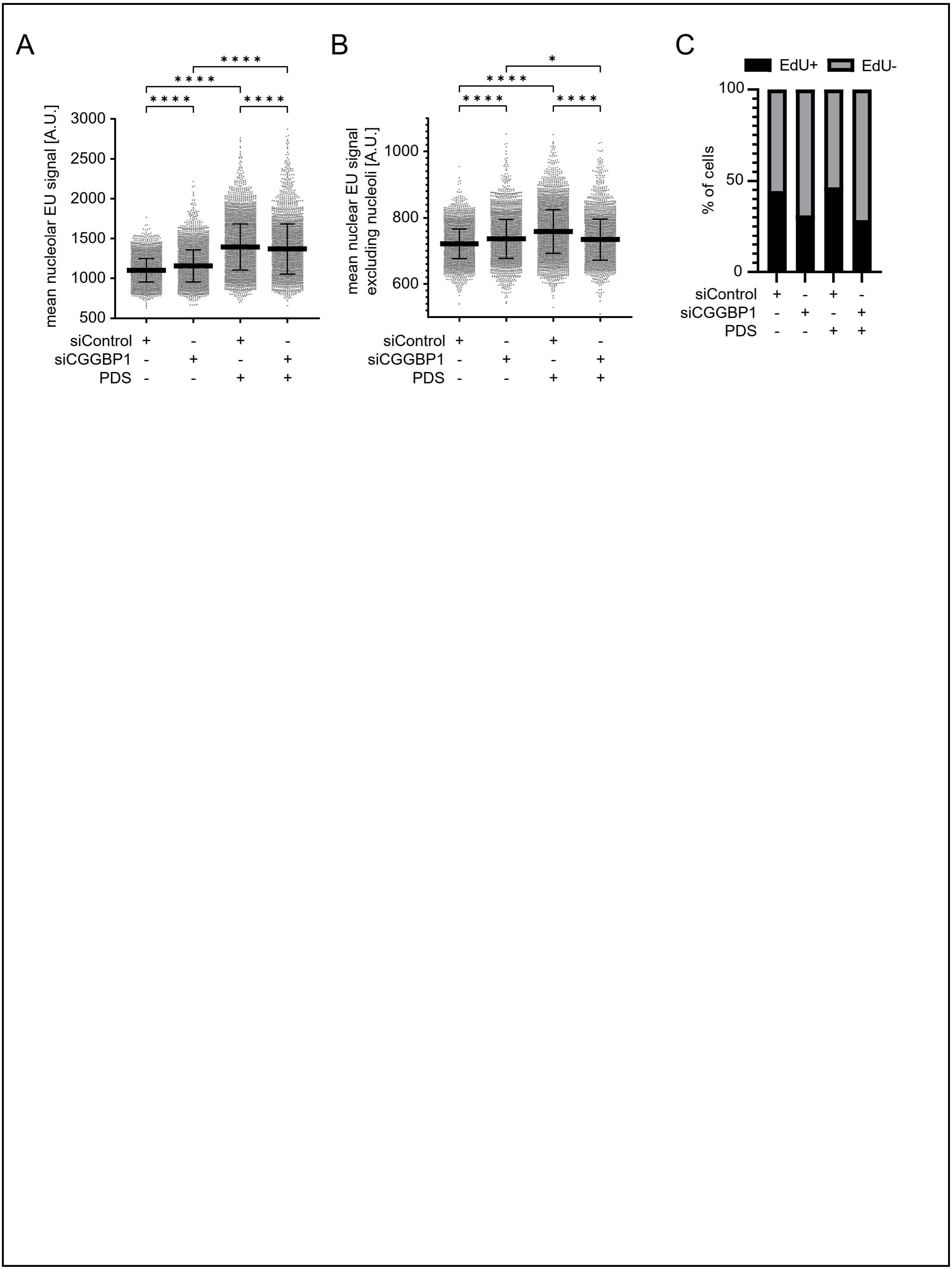

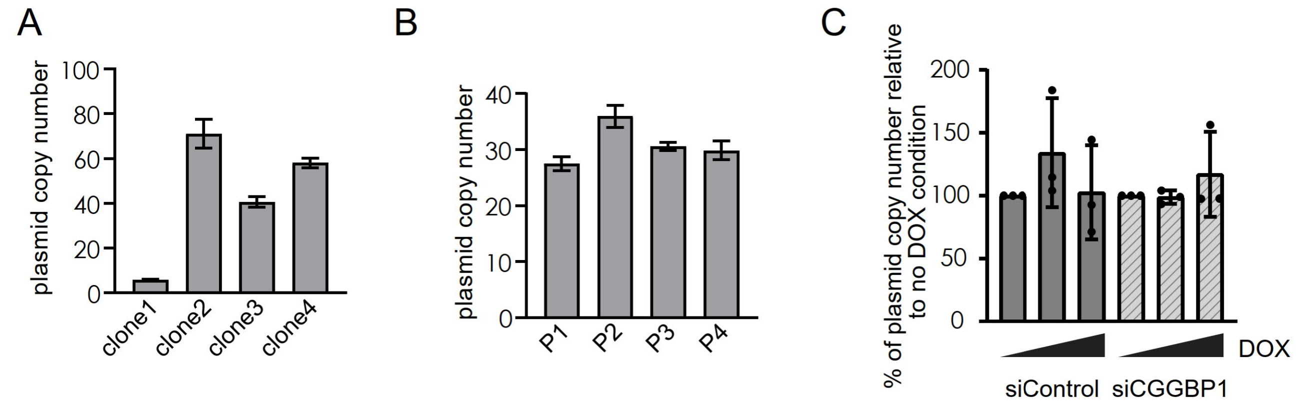
